# Disruption of ribosome dynamics and mRNA homeostasis triggers a cascading imbalance in protein synthesis in models of Amyotrophic Lateral Sclerosis

**DOI:** 10.1101/2025.05.15.654227

**Authors:** Fabio Lauria, Federica Maniscalco, Cecilia Perrucci, Marta Marchioretto, Ilaria Bruno, Gabriele Tomè, Federica Cella, Emma Busarello, Martina Sevegnani, Lorenzo Lunelli, Luisa Donini, Toma Tebaldi, Maria D’Antoni, Daniele Pollini, Daniele Peroni, Alessandra Pisciottani, Laura Croci, Aurora Badaloni, Renato Arnese, Alessandro Provenzani, Alessandro Quattrone, Gian Giacomo Consalez, Massimiliano Clamer, Velia Siciliano, Manuela Basso, Gabriella Viero

## Abstract

The RNA-binding protein TDP-43 is a major contributor and a pathological hallmark of Amyotrophic Lateral Sclerosis (ALS), yet how TDP-43 dysregulation mechanistically alters protein synthesis across neuronal compartments and disease models remains unclear. Here, we dissected TDP-43–driven translational alterations in both in vitro and in vivo TDP-43 models of ALS. Through ribosome and polysome profiling, computational, and biochemical analyses, we observed robust TDP-43-associated translational remodelling at cellular and subcellular resolution. Our findings reveal a conserved mechanism across models, characterized by enhanced ribosome recruitment on polysomes, elongation impairment, axonal downregulation and instability of TDP-43 target mRNAs and redistribution toward non-target transcripts. Notably, TDP-43 dysregulation alters ribosome dynamics and selectively impairs translation of TDP-43 target mRNAs, whilst favouring the translation of other transcripts. This process reflects a compensatory but maladaptive response to TDP-43–induced mRNA destabilization. Together, these data demonstrate that alterations in TDP-43 disrupts neuronal proteostasis through ribosome reorganization and loss of mRNA homeostasis, providing a unifying mechanistic framework for translational imbalance in ALS and revealing early molecular events that may underlie motor neuron vulnerability.

## Introduction

Amyotrophic Lateral Sclerosis (ALS) is a fatal neurological disorder primarily characterised by progressive degeneration of upper and lower motor neurons and alterations in axons and presynaptic terminals in the corticobulbar/corticospinal tract and in the motor component of peripheral nerves (Ito et al., 2017). Around 90% of ALS cases are sporadic, and the remaining 5% have an established genetic cause (Kim et al., 2020; Taylor et al., 2016). Even if our understanding of the genetic basis of ALS has steadily advanced, progress in developing new therapies remains hindered by our limited knowledge of the fundamental cellular and molecular events in affected cells. ALS, as many other neurodegenerative diseases, is characterised by defects in mRNA metabolism, affecting RNA granule formation and translation (Wang and Sun, 2023).

Mutations in RNA Binding Proteins (RBPs) such as TDP-43 (Kabashi et al.2008) and FUS (Kwiatkowski et al.2009; Vance et al. 2009) are major contributors to the development of genetic ALS, most likely by altering mRNA localization, maturation, and translation. TDP-43, which is encoded by the *TARBDP* gene (Buratti et al., 2001), is generally localized to the nucleus, where it is involved in mRNA splicing (Fiesel et al., 2012). In the cytosol and neurites, TDP-43 is involved in RNA transport and granule formation (Corbet et al., 2021; Maharana et al., 2018), binding several thousands of mRNAs in their 3’ UTR through GU repeats (Tollervey et al., 2011; Polymenidou et al., 2011). TDP-43 mislocalisation produces cytoplasmic aggregates leading to the so called TDP-43 proteinopathy (Scotter et al., 2015; Gao et al., 2018). These aggregates are common hallmarks of >90% of all ALS cases (Neumann et al., 2006; Peng et al., 2020). Cytoplasmic accumulation of TDP-43 is associated with broad defects in RNA metabolism, from mRNA mis-splicing (Fiesel et al., 2012), cryptic polyadenylation (Bryce-Smith et al., 2025; Zeng et al, 2025) and defective transport (Alami et al., 2014; Bilsland et al., 2010), to impaired stress granule formation (Colombrita et al., 2009), mRNA degradation (Tank et al., 2018) and altered protein synthesis (Arai et al., 2006; Neumann et al., 2006; Geser et al., 2008; McCluskey et al., 2009; Ling et al., 2013, Arnold et al., 2013; Russo et al., 2017).

Although TDP-43 dysfunction has been shown to cause translation alterations (Piol et al., 2023) either at global or mRNA-specific levels both in *in vitro* (Fiesel et al., 2012; Charif et al., 2020; Neelegadan et al., 2019; Russo et al., 2017; Chu et al., 2019; Altman et al., 2021; Majumder et al., 2012; Blokhuis et al., 2016; Sun et al., 2015) and *in vivo* models of ALS (Gao et al., 2021; Coyne et al., 2015; Lehmkuhl et al., 2021, Charif et al., 2020), no conclusions as to how TDP-43 dysregulation may impact on protein synthesis has been reached. In fact, global protein synthesis has been reported to either increase (Gao et al., 2021, Fiesel et al., 2012, Charif et al., 2020) or decrease in various models of TDP-43 proteinopathy (Gao et al., 2021, Altman et al., 2021, Russo et al., 2017), also within the axon (Pisciottani et al., 2023, Briese et al., 2020). Similarly, transcript-specific translational defects were observed (Coyne et al., 2014, Chu et al., 2019; Majumder et al., 2016, Majumder et al., 2012), showing the complex alterations in protein synthesis in multiple models of ALS (Bjork et al., 2022, Wang and Sun, 2023).

To dissect the translational alterations driven by TDP-43 proteinopathy across neuronal compartments and disease models, we employed primary cortical neurons overexpressing TDP43 mutants (Buratti, 2015) cultured in microfluidic chambers enabling compartmentalized analysis of cell bodies and axons (Pisciottani et al., 2023). We integrated multiple complementary approaches, including ribosome and polysome profiling, RNA-Seq, and biochemical assays, and systematically examined the interplay between RNA granules and polysomes also at subcellular resolution. Comparative analyses across an *in vivo* mouse model of ALS (Arnold et al., 2013) and publicly available RNA-Seq datasets from ALS patient post-mortem spinal cords (Krach et al., 2018) revealed a conserved, disease-model-independent mechanism characterized by axonal downregulation of translation for TDP-43 targets and a reorganization of ribosome recruitment on neuronal mRNAs. Together, these data uncover a shared translational remodelling program underlying TDP-43–mediated neurodegeneration.

## Results

### Ribosome reorganization into larger polysomes is a common feature in ALS models

To gain mechanistic insight into translational alterations driven by TDP-43 proteinopathy, we first quantitatively assessed its association with the translation machinery, including ribosomal subunits, ribosomes, polysomes, and RNA granules. Although prior studies suggested that TDP-43 may interact with polysomes (Freibaum et al., 2010; Higashi et al., 2013; Coyne et al., 2015; Russo et al., 2017; Coyne et al., 2014; Majumder et al., 2016; Neelagandan et al., 2019), a systematic, quantitative analysis across cell types and tissues has been lacking. Using polysome profiling in human and mouse cell lines and mouse tissues (**Fig. 1a**), we isolated proteins and RNAs associated with ribosomes/polysomes or with ribosome-free “ribonucleoprotein particles (RNPs)/granules”. Using established markers of the small (RPS6) and large (RPL26) ribosomal subunits and of RNA granules (FUS and HuD) (**Supplementary Fig. 1a-c** and **Supplementary File 1**), we found that under physiological conditions, most of the endogenous TDP-43 protein co-sediments with RNPs/granules. This pattern was consistent in human induced pluripotent stem cells (hiPSCs), murine motor neuron-like cell lines (NSC-34) and murine cortex and spinal cord (**Fig.1b** and **Supplementary Fig. 1a-c**). Notably, a small fraction of the total TDP-43, ranging between 2% and 5%, was found to associate with the translation machinery (**Fig. 1b**). To understand whether the association of endogenous TDP-43 with the translation machinery is exclusively RNA-dependent, we performed a subcellular fractionation assay in hiPSCs with and without RNAse treatment (**Fig. 1c** and **Supplementary File 2**), as described in Lauria et al., 2020. The ribosomal proteins of the small (RPS6) and large (RPL26) subunits were used as markers of ribosomes and polysomes, FUS as a marker of RNA granules, and the polyA-binding protein (PABP) as a control of mRNA-dependent association of proteins with polysomes. In agreement with what was observed in **Fig. 1b** (left donut plot) and **Supplementary Fig. 1a,** in the absence of RNAse treatment (**Fig. 1c**, left panel), the large majority of TDP-43 primarily associates with RNPs/granules (S100 lane). After a step of high salt wash, the association of TDP-43 with the ribosome pellet is still visible (WR lane), suggesting that TDP-43 directly binds the translation machinery. As suggested by its binding with RACK1 (Russo et al., 2017), after endonuclease digestion (**Fig. 1c**, right panel), TDP-43 is still associated with ribosomes (R and WR lanes) through RNA-independent interactions.

**Fig. 1.**
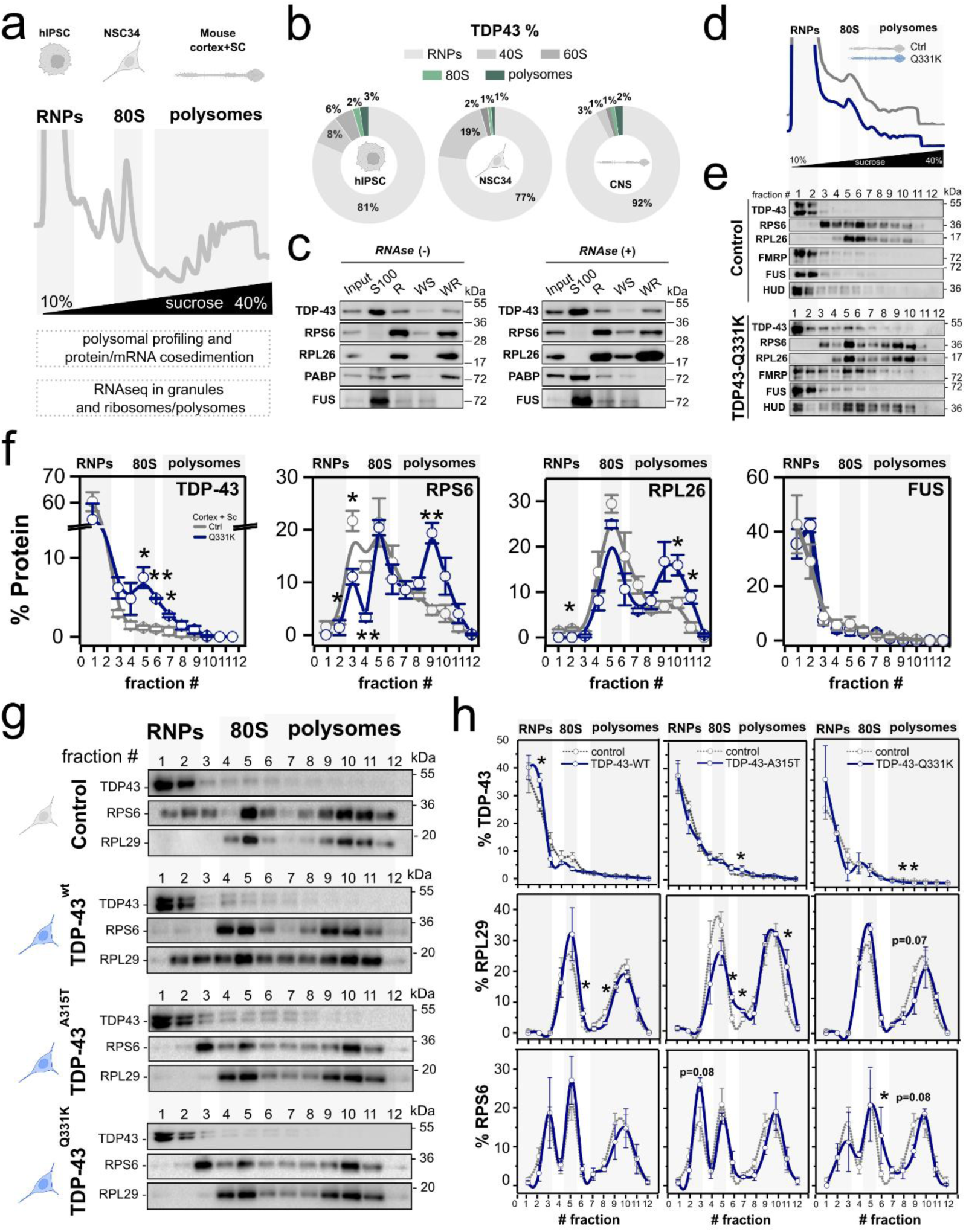
Ribosome reorganization into larger polysomes is a common feature in ALS models. (**a**) Schematic representation of the biological systems and approaches used in this paper to investigate TDP-43’s role in translation. (**b**) Donut plots reporting the relative percentages of endogenous TDP-43 co-sedimentation with RNPs, 40S, 60S, 80S, and polysomes in polysome profiles of hiPSCs, NSC-34 cell lines, and murine central nervous system (CNS). Data are mean of n=3 biologically independent replicates. (**c**) Western blot analysis of subcellular fractions before (left) and after (right) RNAse I treatment of cytoplasmic lysates from hiPSCs (input). Labels correspond to ribosome-free cytoplasmic components (S100); ribosomes and polysomes (R); loosely ribosome-bound proteins (WS); and strongly ribosome-bound proteins (WR). Ribosomal markers RPL26 and RPS6 were used as control of ribosome sedimentation. FUS was used as a marker of RNA granules. PABP was used as control for RNA-dependent interactions. (**d**-**e**) Representative polysome profiles (d) and representative co-sedimentation profiles (e) of TDP-43, ribosomal markers (RPL26, RPS6), and RNA-binding proteins (FMRP, FUS, and HUD) in the cortex and spinal cord of control and transgenic mice expressing hTDP-43-Q331K. (**f**) Relative percentage distribution along the profile of TDP-43, ribosomal markers (RPS6, RPL26), and FUS in the cortex and spinal cord of control and transgenic mice expressing hTDP-43-Q331K. Data are mean ± SEM of n=3 biologically independent replicates. Differences were assessed with two-tailed T-test (P-value: *<0.05, **<0.01). (**g**) Representative co-sedimentation profiles of TDP-43 (s: short exposure; l: long exposure; upper bands: His-HA-tagged hTDP-43-WT, lower bands: endogenous TDP-43-WT) and ribosomal markers (RPL29, RPS6) in NSC-34 expressing His-HA-tagged hTDP-43-WT or two hTDP-43 mutants (A315T and Q331K). (**h**) Relative percentage distribution along the profile of TDP-43 and ribosomal markers (RPL29, RPS6) in NSC-34 expressing His-HA-tagged hTDP-43-WT or two hTDP-43 mutants (A315T and Q331K). Data are mean ± SEM of n=3 biologically independent replicates. Differences were assessed with two-tailed T-test (P-value: *<0.05, **<0.01). hIPSC, NSC34 and mouse cortex+SC illustrations created in BioRender.

Next, we explored the hypothesis that TDP-43 mutations, which are involved in TDP-43 proteinopathy (Buratti, 2015; Scotter et al., 2015), change the balance between TDP-43 association with RNPs/granules and the translation machinery. Since mild overexpression of WT or mutant TDP-43 in neurons leads to TDP-43 proteinopathy and ALS-like neuronal phenotypes (Tsai et al., 2010), we used the following as models of ALS: i) cortices and spinal cords of an *in vivo* murine model of ALS characterised by the transgenic expression in the central nervous system of the mutant human TDP-43-Q331K (hTDP-43-Q331K) (Arnold et al., 2013) (**Supplementary Fig. 1d** and **Supplementary File 1**); ii) three murine motor neuron-like cell lines (NSC-34) stably engineered for the doxycycline-inducible expression of His-HA-tagged hTDP-43-WT or two hTDP-43 mutants (A315T and Q331K) (**Supplementary Fig. 1e** and **Supplementary File 1**). After polysome profiling, we observed no changes in ribosomes in polysomes (FRP) (**Fig. 1d** and **Supplementary Fig. 1f,g)**, consistent with previous data in MN1 cell lines (Neelagandan et al., 2019). However, we observed that hTDP-43-Q331K significantly increases its association with ribosomes and lighter polysomes (**Fig. 1e,f**), in the cortex and spinal cord of the *in vivo* model of ALS at an early symptomatic stage of the disease (**Fig. 1e-h** and **Supplementary File 2**). No changes were observed for FUS protein, another RBP involved in ALS (**Supplementary Fig. 1f,g**). The change in hTDP-43-Q331K is accompanied by a redistribution of the ribosomal markers RPL26 (large subunit) and RPS6 (small subunit), which shifts towards polysomes with a higher number of ribosomes (**Fig. 1e,f**). In agreement with these results, in murine motor neuron-like models of TDP-43 proteinopathy (**Fig. 1g,h**) we found that both TDP-43-WT and TDP-43-Q331K accumulated in the RNPs/granule fractions, whilst TDP-43-A315T exhibited an increased association with polysomes (**Fig. 1g,h**).

Together, these findings indicate that, despite milder model-specific differences, TDP-43 association with the translation machinery increases in the mutant background, promoting a consistent shift of ribosomal subunits toward high-number polysomes.

### Mutant TDP-43 induces a compartment-specific rewiring of neuronal translation

As TDP-43 proteinopathy mostly affects neurons with long projections, we wondered if the observed redistribution of ribosomes occurs preferentially in axons or cell bodies. To test this, we cultured Primary Cortical Neurons (PCNs) in microfluidic chambers and applied a miniaturized polysome profiling strategy optimized for low-input samples to generate sedimentation profiles (**Fig. 2a**, **Supplementary Fig. 2a,b**). After validating that ribosomal co-sedimentation patterns were consistent with traditional sucrose gradients (Lauria et al., 2020) (**Supplementary Fig. 1c**; **Supplementary File 3**), we analyzed the sedimentation of ribosomes in neurons expressing either the control Turbo Red Fluorescent Protein (TRFP) or TDP-43-A315T–TRFP (**Fig. 2b,c**; **Supplementary Fig. 2d–h**; **Supplementary Files 3,4**). The distribution of the ribosomal markers in control samples helped us identify the ribosome and polysome fractions, which in the control cell body are in fractions 7-8 and 9-12, respectively (**Figure 2b**, left panels). In agreement with what shown in whole cells and tissues, in the cell bodies the endogenous TDP-43 is primarily associated with RNPs/granules, whilst in the axons the polysome-associated endogenous TDP-43 accounts for 1/5 of the axonal TDP-43 (**Fig. 2d)**. In agreement with what observed in NSC-34 cells expressing TDP-43-A315T (**Fig. 1g,h**), we found that the association of TDP-43-A315T with polysomes increases in the cell body (**Fig. 2e**, left panels). Conversely, in the axon, we did not observe alterations in the association of TDP-43 with ribosomes or polysomes but a slight shift of the ribosomal proteins towards RNPs/granules fractions (**Fig. 2e**, right panels).

**Figure 2.**
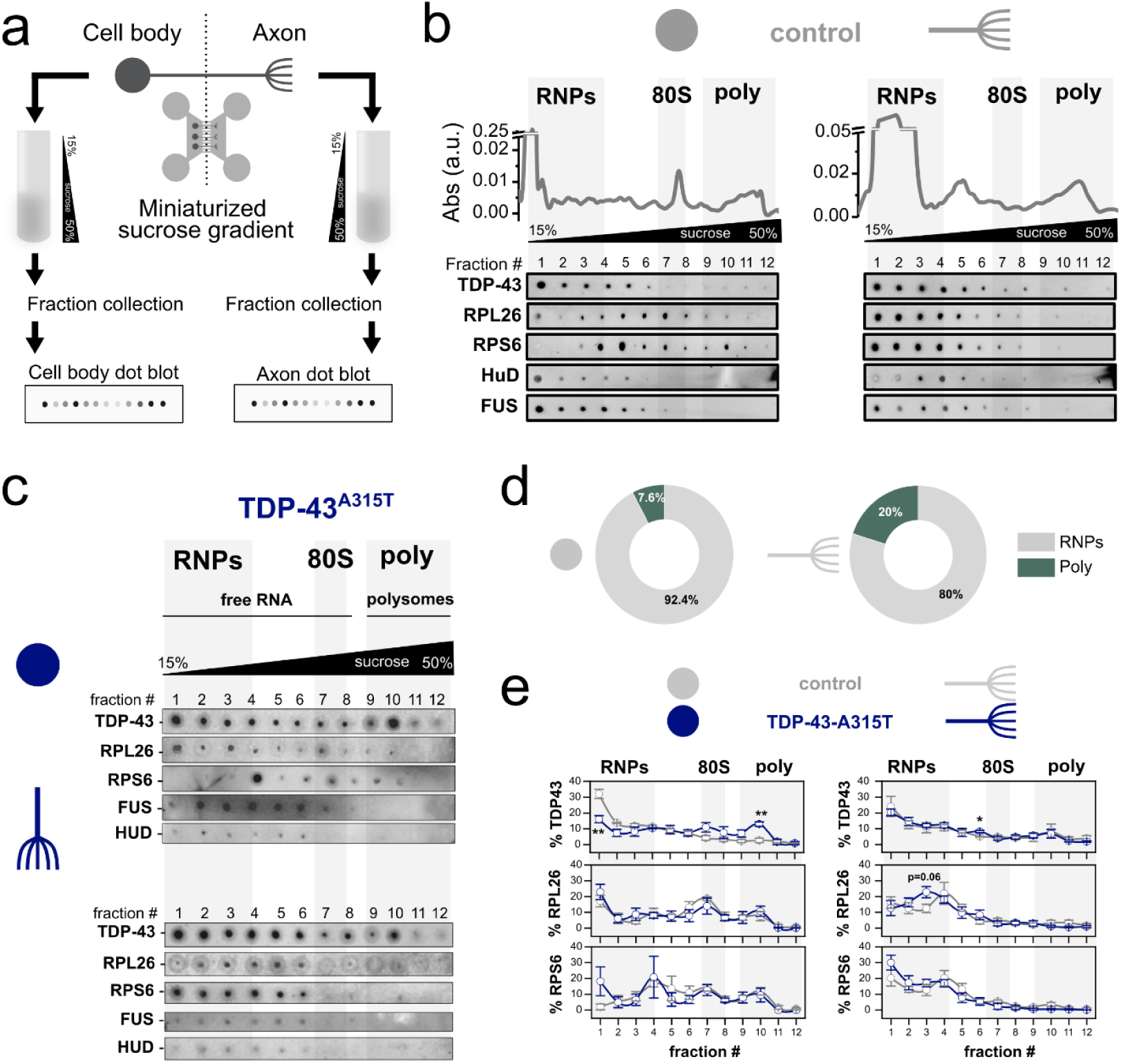
Mutant TDP-43 induces a compartment-specific rewiring of neuronal translation. (**a**) Experimental design for the isolation of cell bodies and axon from cortical neurons cultured in microfluidic chambers. . (**b-c**) Representative polysomal profiles and dot blot co-sedimentation profiles of TDP-43, ribosomal markers (Rpl26 and Rps6), and RNA-binding proteins (FMRP, Fus and Hud) in cell body and axons from wild type Primary Cortical Neurons (PCNs) (c) and PCNs upon ectopic expression of TDP-43-A315T (d) cultured in microfluidic chambers. (**d**) Donut plots with relative percentages of endogenous TDP-43 association in RNPs/granules and polysome compartments in the cell body (left panel) and axons (right panel) of wild type PCNs. (**e**) Quantification of co-sedimentation profiles of TDP-43 and ribosomal markers in the cell body and axons of PCNs control and expressing TRFP-TDP-43-A315T. Data are mean ± SEM of n=3-4 biologically independent replicates. Differences were assessed with two-tailed T-test (P-value: *<0.5, **<0.1).

These findings reveal that TDP-43-A315T selectively enhances polysome association in the cell body while subtly redirecting ribosomal proteins toward RNPs/granules in axons, pointing at a compartment-specific rewiring of neuronal translation.

### TDP-43 proteinopathy alters occupancy of TDP-43 target mRNAs at early stages of disease

Since TDP-43 proteinopathies drives a consistent shift of ribosomal subunits toward polysomes, we speculated that these changes might affect their occupancy on mRNAs. To test this hypothesis, we performed traditional ribosome profiling (Ingolia et al., 2009) and in parallel RiboLace (Clamer et al., 2018) (**Fig. 3a**). The first method helps identify both active and inactive (or stalled) ribosomes, while the second specifically provides ribosome occupancy data for active ribosomes. Although ribosome positioning at the start and end of mRNAs was unchanged (**Supplementary Fig. 3a**), we detected widespread alterations in ribosome occupancy (**Fig. 3b**, **Supplementary Fig. 3b**; **Supplementary File 5**) that were largely independent of active ribosome levels (**Fig. 3b,c**). Enrichment analysis revealed that transcripts with reduced ribosome occupancy are preferentially bound by TDP-43 (**Fig. 3d**), indicating that ribosome alterations in this ALS model predominantly affect TDP-43 target mRNAs.

**Figure 3.**
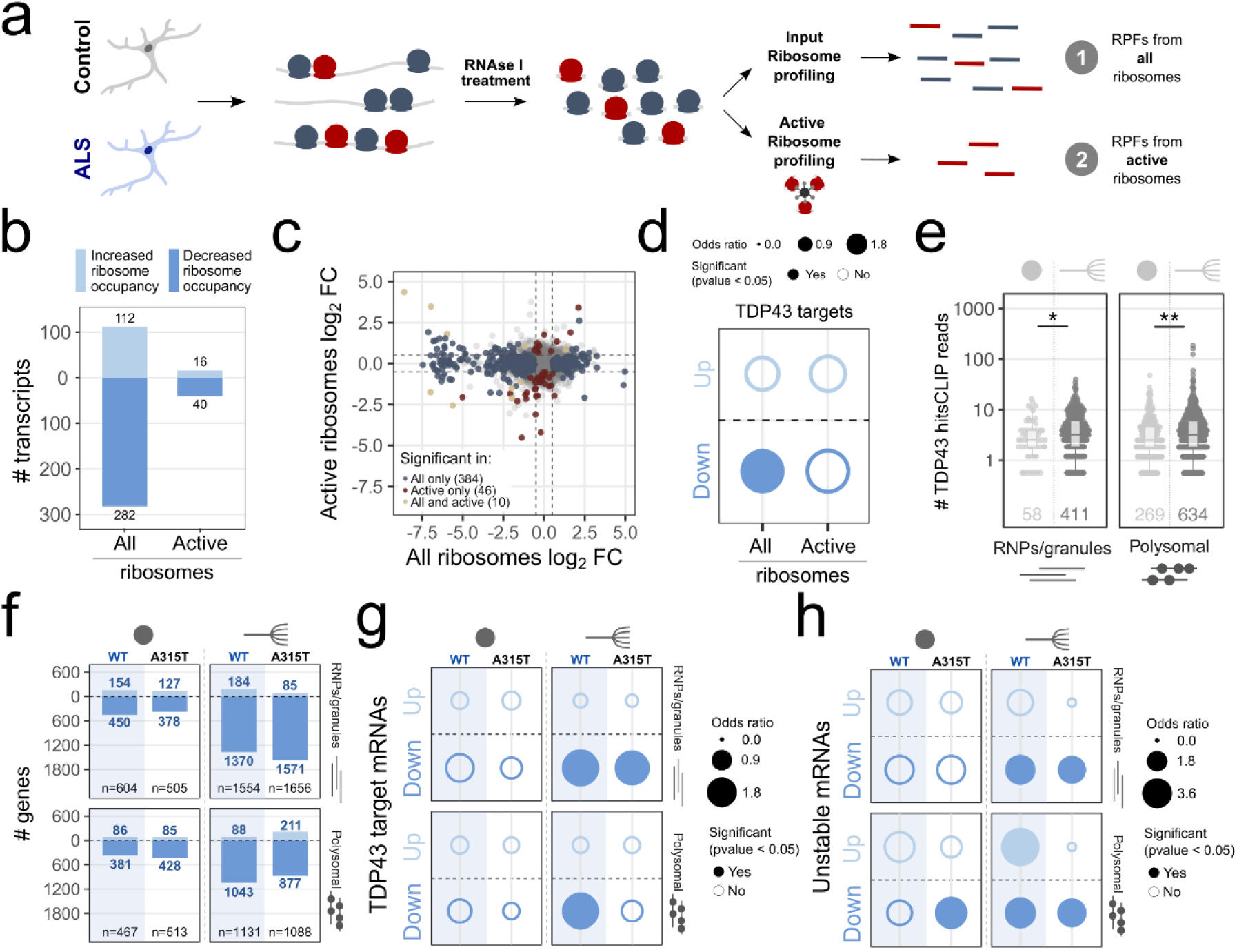
TDP-43 proteinopathy alters occupancy of TDP-43 target mRNAs at early stages of disease. (**a**) Schematic of the experimental design for studying the localization of all ribosomes (either active or inactive) and only active ribosomes. (**b**) Number of mRNAs showing decreased or increased ribosome occupancy in ALS considering both active and inactive ribosomes (“All”) or only active ribosomes. (**c**) Comparison of ribosome occupancy variations based on both active and inactive ribosomes (“All”, x-axis) and on active ribosomes (y-axis). The colour reports whether the variation is significant only if considering both active and inactive ribosomes (“All only”), only according to the active ribosomes (“Active only”) or both populations (“All and active”). (**d**) Enrichment analysis of TDP-43 target mRNAs among transcripts showing increased and decreased ribosome occupancy in ALS. Dot size is proportional to the enrichment measured by odds ratio and the significant enrichments are reported as filled circles (Fisher’s exact test). (**e**) Number of TDP-43 CLIP-Seq hits (Polymenidou et al., 2011) for polysomal and RNPs/granules enriched in the cell body chamber and in axons of PCN expressing TRFP. The number of genes is shown at the bottom of the boxes. Differences were assessed with two- two-tailed T-test (P-value: * < 0.05, ** < 0.01). (**f**) Bar plot showing the number of up- and down-regulated mRNAs in the cell body and axon upon TDP-43-WT and TDP-43-A315T expression with respect to TRFP. The number of DEGs for each comparison is reported next to the bars and the total number of DEGs is shown at the bottom of the boxes. (**g-h**) Enrichment analysis of TDP-43 target genes (g) and of destabilised genes due to the expression of TDP-43 (Tank et al., 2018) (h) among up- and down-regulated genes identified upon TDP-43-WT or TDP-43-A315T expression. Dot size is proportional to the enrichment measured by odds ratio and the significant enrichments are reported as filled circles (Fisher’s exact test).

To gain subcellular resolution, we determined the catalogue of RNP/granule-associated RNAs and polysome-associated RNAs from both cell bodies and axons of PCNs cultured in microfluidic chambers by RNA-Seq ALS expressing TDP-43-WT or TDP-43-A315 (Pisciottani et al., 2023) (**Supplementary Fig. 3c-e**). The expression of Map2 and Nfl mRNAs, as markers of dendrites and neurons, respectively (Briese et al., 2016), showed no dendritic contaminations in the axonal compartment (**Supplementary Fig. 3f** and **Supplementary File 6**). Importantly, mRNAs in the control axonal RNPs/granule and polysome compartments are more likely to be bound by TDP-43 than mRNAs localised in the cell body (**Fig. 3e**), suggesting that mRNAs associated with axonal RNPs/granules and axonal polysomes may be more sensitive to TDP-43 proteinopathy. The differential expression analysis at the cell body level revealed 1071 (TDP-43-WT) and 1018 (TDP-43-A315T) differentially expressed genes (DEGs) (**Fig. 3f**). Of these DEGs only 58 and 72 are shared across the RNPs/granules and polysomal compartment, respectively. Strikingly, in the axon the number of DEGs is almost three times larger than in the cell body (2685 for TDP-43-WT and 2744 for TDP-43-A315T), accounting for 89% of the total changes (**Fig. 3f** and **Supplementary File 7**). The number of DEGs shared across the RNPs/granules and polysomal compartment, are ∼ 1/10, i.e. 278 (TDP-43-WT) and 213 (TDP-43-A315T). Interaction network analysis of proteins encoded by axonal DEGs in both conditions revealed shared (**Supplementary Fig. 3g**) or specific (**Supplementary Fig. 3h**) protein communities. Interestingly, the emerging communities are mainly associated with localization/transport, catabolic and metabolic processes, cell projection, and mitochondrion (**Supplementary Fig. 3g-h**). These communities recall several themes known to be altered in ALS (Briese et al., 2020, Nagano et al., 2020, Lehmkul et al., 2021, Guise et el., 2024; Pisciottani et al., 2023). As in the ribosome profiling analysis (**Fig. 3d**), we observed a significant enrichment of TDP-43 targets in the downregulated set of DEGs (**Fig. 3g**). This enrichment is axonal-specific (**Fig. 3g**, right panels), as no enrichment was associated with the cell body compartment (**Fig. 3g**, left panels) or with up-regulated genes. Consistent with the role of TDP-43 in RNA destabilization (Ayala et al., 2011; Tank et al., 2018), downregulated axonal targets are enriched in unstable RNAs, while upregulated genes are not (**Fig. 3h**)

While most neurodegenerative disorders are late onset, it is possible to identify very early and translation specific signatures of the disease (Bernabò et al., 2017, Genovese et al., 2025). To test if also TDP-43 proteinopathy cause *in vivo* and early-symptomatic alterations in ribosome dynamics at very early stages of the disease, we shortlisted 21 genes, selected as follows: i) 11 DEGs shared between our results and a dataset from ALS patients (Krach et al., 2018), showing altered ribosome occupancy in human motor neurons after TDP-43 silencing (Brown et al., 2022) (**Fig. 4a** and **Table 1**), ii) 5 DEGs shared between our results and the dataset from patients (Krach et al., 2018), iii) 5 genes not differentially expressed genes. Among the selected genes, 13 genes are TDP-43 target mRNAs and the remaining 8 are not. We used Actin-b as a housekeeping gene because it shows no variations either at mRNA and protein levels (**Supplementary Fig. 4a** and **Supplementary File 8**). In the spinal cord and cortex of ALS early symptomatic mice, we observed a trend of transcriptional downregulation (**Fig. 4b**), as in our previous analysis. Next, we analysed the association of the 21 transcripts with polysomes using the relative distribution of the mRNAs with the translation machinery in WT and ALS mice (**Fig. 4c** and **Supplementary Fig. 4b,c**). We observed that in the mutant background TDP-43 non-target mRNAs increase their association with the 80S/monosome fraction, while TDP-43 target mRNAs significantly shift toward the heavier polysome fraction (**Fig. 4d**).

**Table 1.**

| Gene | Description | References |
| --- | --- | --- |
| C1orf216 | chromosome 1 open reading frame 216 |  |
| CTDSP2 | CTD small phosphatase 2 |  |
| EGR1 | early growth response 1 | Ledener et al., 2007; Moujalled et al. 2017; Godini et al., 2018 |
| GSTM4 | glutathione S-transferase mu 4 | Klim et al., 2019; Cha et al., 2020 |
| HES1 | es family bHLH transcription factor 1 | von Grabowiecki et al., 2015; Yang et al., 2015 |
| PSAT1 | phosphoserine aminotransferase 1 | Martins De-Sousa et al., 2012 |
| PTPRT | protein tyrosine phosphatase receptor type T | Carter et al., 2009; Liu et al. 2020 |
| RRBP1 | ribosome binding protein 1 | Tank et al., 2018 |
| SLC1A5 | solute carrier family 1 member 5 | Lee et al., 2016 |
| SLC7A11 | solute carrier family 7 member 11 | Vallerga et al., 2020 |
| VIM | Vimentin | Zhou et al., 2015 |

**Figure 4.**
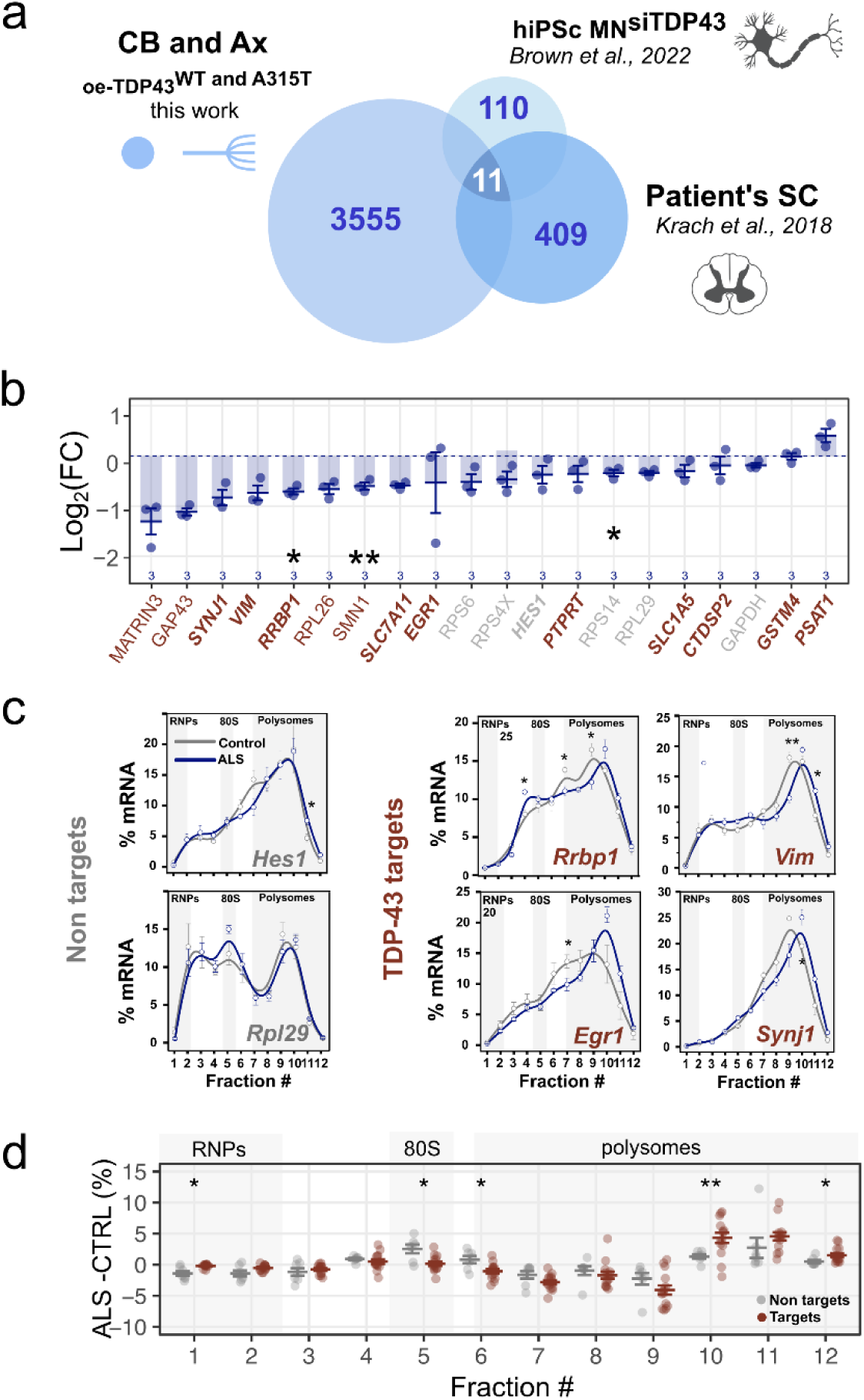
TDP-43 targets show translational defects at an early symptomatic stage of the disease. (**a**) Overlap between differentially expressed genes (DEGs) identified in our *in vitro* model of ALS, in spinal cords of ALS patients (Krach et al., 2018) and hiPSc motor neurons (Brown et al., 2022), compared to the control condition. Only genes expressed in all samples were considered. The 11 shared genes are listed in Table 1. MN and SC illustrations created in BioRender. (**b**) qPCR-derived fold changes in mRNAs for selected TDP-43 targets (red text) and non-target (grey text) genes upon expression of hTDP-43-Q331K in mouse spinal cords and cortices. Data were normalised to Actin-b. The number of biologically independent replicates is reported at the bottom of the box. Data are mean ± SEM. Differences were assessed with two-tailed T-test (P-value: *<0.05, **<0.01). (**c**) Representative relative co-sedimentation profile of TDP-43 non-targets mRNAs and weak, medium and strong TDP-43 targets mRNAs along the sucrose gradient fractions of control and ALS mouse spinal cords and cortices. Data are mean ± SEM of n=3 biologically independent replicates. Differences were assessed with two-tailed T-test (P-value: *<0.05, **<0.01). (**d**) Differences between ALS and control relative percentage co-sedimentation along the sucrose gradient fractions. Data are mean ± SEM of 8 TDP-43 non-target and 12 TDP-43 target genes. Differences were assessed with two-tailed T-test (P-value: *< 0.05, **<0.01).

Thus, our data show how TDP-43 pathology is characterized by a differences between non target and TDP-43 target mRNAs, which show a widespread destabilization and a shift to polysome fractions already at an early symptomatic stage.

### TDP-43 rewires axonal polysome–RNP partitioning of target mRNAs and translational efficiency

The widespread instability of axon-enriched TDP-43 targets likely limits their ribosome availability, thereby causing systemic rewiring of translation and broad effects on gene expression. This concept reflects cellular resource reallocation, the process by which a cell reallocates its resources, i.e. ribosomes in this case, toward different processes to adapt to shifts in cellular conditions and resource abundances (Frei et al., 2020).

To first get an insight on global translation dynamics at the axonal level, we used an *in situ* puromycin-based labelling approach in cortical neurons (David et al., 2012) and found an increase in the number of axonal foci of active translation in the axons of PCNs overexpressing TRFP-TDP-43-WT and TRFP-TDP-43-A315T (**Figure 5a**).

**Figure 5.**
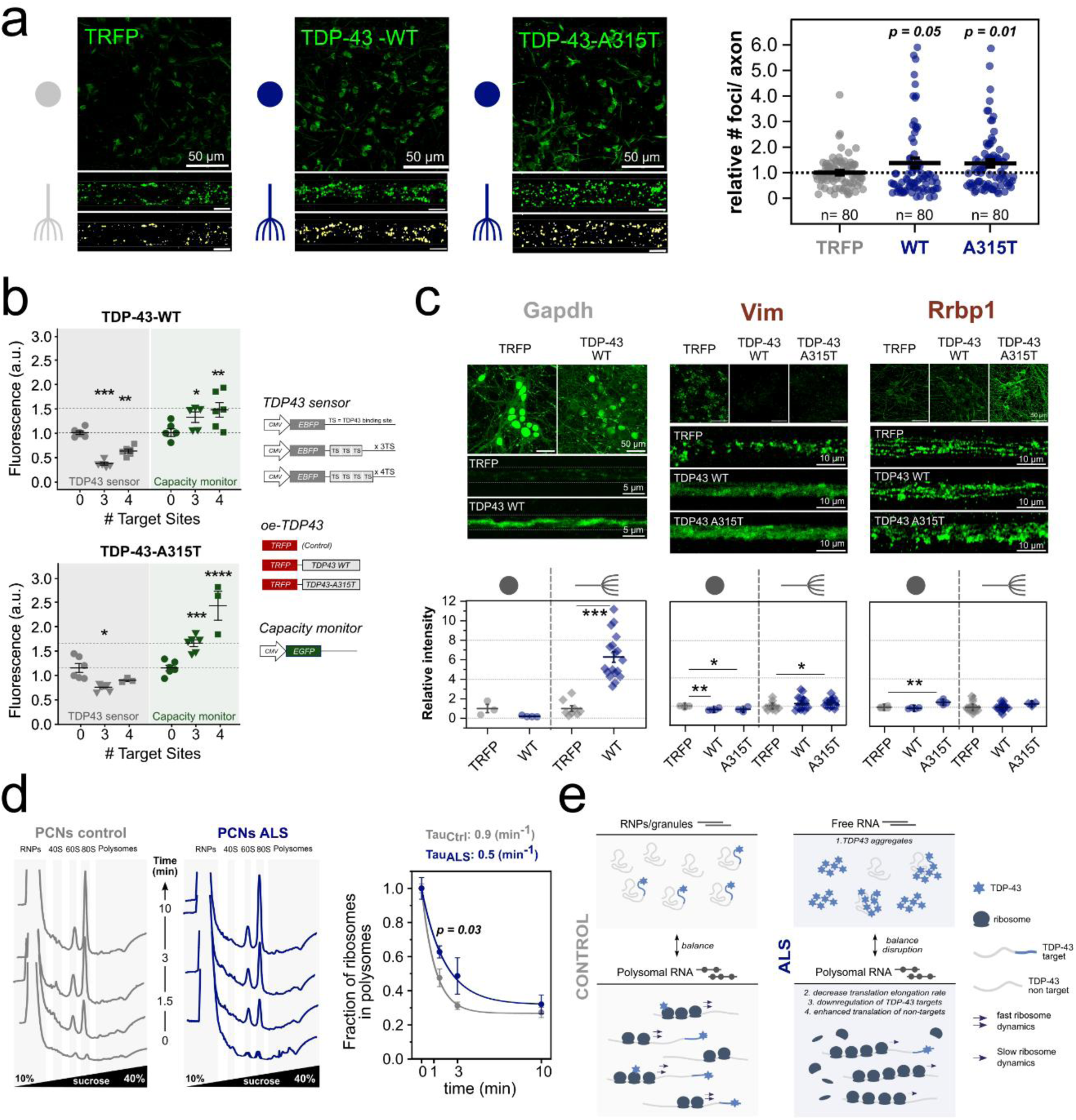
TDP-43 rewires axonal polysome–RNP partitioning of target mRNAs and translational efficiency. (**a**) Left panels: Immunofluorescence of cell bodies and axons in microgrooves of microfluidic chambers after puromycilation assay. Neurons infected with TRFP, TDP-43-WT, TDP-43-A315T were treated with puromycin, and sites of active translation were labelled with Alexa 488 nm (green dots). Right panel: average number of foci of active translation per 10 μm axon length in axons of cortical neurons expressing TDP-43-WT or TDP-43-A315T relative to TRFP controls. Differences were assessed with two-tailed T-test. P-values are reported in the figure. (**b**) Left panels: relative fluorescence signal of TDP-43 sensor (gray) and capacity monitor (green) upon TDP-43-WT and TDP-43-A315T expression, respectively. Data are mean ± SEM of n=3-6 biologically independent replicates. Differences were assessed with two-tailed T-test (P-value: *<0.05; **<0.01; ***<0.001; ****<0.0001). Right panels: schematic representation of plasmids used in the translational burden assay where NSC-34 cells were co-transfected with 3 plasmids to monitor TDP-43 target and non-target translation in control and diseased condition. Bottom panels: (**c**) Representative images of immunofluorescence (upper panels) and relative fluorescence intensity quantification (bottom panels) of one non-target (Gapdh) and two TDP-43 targets (Vim and Rrbp1) in the cell body and in axons of PCNs cultured in microfluidic chambers and infected with TRFP control, TDP-43-WT or A315T. Differences were assessed with two-tailed T-test (P-value: *<0.05, **<0.01, ***<0.001). (**d**) The effect of harringtonine on polysome profiles from control PCNs (left panel) and PCNs expressing hTDP-43-Q331K (middle panel) treated only with cycloheximide (time point 0) and after 1.5, 3, and 10 minutes of harringtonine treatment. Right panel: FRP calculated from the polysome profiles of harringtonine treated cells for each time point: data were fitted with an exponential decay curve to compute the ribosome kinetics (Tau) values, which is 1.8 times slower in ALS (tALS= 0.5 min-1) compared to control (tCTRL= 0.9 min-1) condition. Data are mean SEM of n=5-7 biologically independent replicates. Differences were assessed with two-tailed T-test (P-value: * < 0.05) (**e**) Proposed working hypothesis of translational effects of TDP-43 overexpression.

As our data on TDP-43 target point at their instability, whilst axonal translation foci are increased, we measured the effect of TDP-43-WT or TDP-43-A315T on TDP-43 targets vs non-targets mRNA (**Fig. 5b**). To this end, we developed a fluorescent sensor including 3 or 4 TDP-43 binding sites (EBFP-TS, namely TDP-43 sensor) to report for TDP-43 target translation, while expressing at the same time an EGFP lacking any TDP-43 binding site (capacity monitor) (**Fig. 5b**, upper panel). This design is based on the expectation that ribosome reorganization would increase translation of the TDP-43-independent sensor while reducing translation of the TDP-43 sensor. In keeping with our hypothesis, we observed that the higher the number of TDP-43 binding sites in the sensor, the less efficient is its translation upon overexpression of TDP-43-WT or TDP-43-A315T compared to controls (**Fig. 5b**, bottom panel, grey). Conversely, the capacity monitor, mimicking TDP-43 non-target mRNAs, is more efficiently translated in both ALS-like models (**Fig. 5b**, bottom panel, green). Further supporting the hypothesis of cellular resource reallocation, in the axon of PCN of ALS mice we also observed a robust increase in protein production for the non-target RNA Gapdh, whilst the TDP-43 targets Vim and Rrbp1 undergo minor or no significant changes (**Fig. 5c**).

While TDP-43 proteinopathy drives a consistent shift of translation towards TDP-43 non-target mRNA, we have shown that TDP-43 target mRNA co-sediments with heavier polysomes (**Fig. 4d**), making us hypothesize that these polysomes might display impaired translation kinetics. To test this, we determined the ribosome kinetics in PCNs isolated from the mouse model of ALS expressing hTDP-43-A331K and littermate controls, by performing a ribosome run-off assay (**Fig. 5d**). Using polysome profiling, we monitored the kinetics of polysome disappearance upon treatment with harringtonine (**Fig. 5d**, middle panel), which selectively blocks ribosomes at the start codon (Fresno et al., 1977). After deriving the time constants, we found a significant decrease in ribosome kinetics in ALS (**Fig. 5d**, right panel). Stemming from these findings, we propose a model of ALS based on slower ribosome dynamics that may cause a drop in the levels of TDP-43 target mRNAs, release of ribosomes, and their reorganization on the remaining mRNAs, mainly from those not normally regulated by TDP-43 (**Figure 5e**).

## Discussion

Although TDP-43 dysfunction has long been associated with alterations in translation across multiple ALS models (Fiesel et al., 2012; Charif et al., 2020; Neelegandan et al., 2019; Russo et al., 2017; Chu et al., 2019; Altman et al., 2021; Majumder et al., 2012; Gao et al., 2021; Lehmkuhl et al., 2021), the precise mechanisms by which TDP-43 dysregulation affects ribosome organization and protein synthesis have remained unclear. In this study, we moved beyond previous qualitative observations of TDP-43–polysome interactions (Coyne et al., 2014; Majumder et al., 2016; Russo et al., 2017; Neelagandan et al., 2019) to provide a quantitative framework describing how TDP-43 associates with the translational machinery across different cellular systems.

We quantified the fraction of endogenous and cytoplasmic TDP-43 bound to ribosomes and found to be <5%. Strikingly, this association was tenfold higher in primary cortical neurons (PCNs), particularly within axonal polysomes enriched in TDP-43 targets. When TDP-43 mutations cause an increase in its cytoplasmic abundance, this leads to alterations in ribosome engagement with polysomes and, in a subcellular contexts, to axonal mRNAs being disproportionately vulnerable to TDP-43 pathology. Using *in vitro* ALS models and improving the resolution achieved by other omics analyses of cell bodies and axons obtained *in vitro* (Nijssen et al., 2018; Briese et al., 2016; Glock et al., 2021) and *in vivo* (Shigeoka et al., 2016), we found widespread mRNA instability, which is an event shared across multiple models of ALS and not accompanied by a decrease in RNA synthesis (Barmada et al., 2015). These events may be triggered by the loss of TDP-43 function resulting in the inclusion of cryptic exons into mRNAs and Nonsense-Mediated Decay (NMD) (Tan et al., 2016; Tziortzouda et al., 2021; Ling et al., 2015), alternative polyadenylation (Bryce-Smith et al., 2025; Zeng et al, 2025), or by a decrease in translation elongation rates (Duviau et al., 2023; Schoemaker and Green 2012, Dong et al., 2024), which we observed as decrease in ribosome dynamics in PCNs purified from an *in vivo* model of ALS. Since our data on TDP-43 targets are derived from a reporter system that bypasses the processes of splicing and polyadenylation, both of which have been reported to be affected in TDP-43 proteinopathy, we propose an additional effect on target mRNA stability. This effect appears to be independent of nuclear processing of immature mRNA and instead may depend on either a direct interaction with TDP-43 or on the suboptimal translation of these mRNAs.

The robust down-regulation of both RNA/free and polysomal TDP-43 targets in the axon can be explained by the reorganization of ribosomes in the cells. This process can be framed within the concept of reallocation of cellular resources (Frei et al., 2020; Cella et al., 2023), i.e. mRNAs and ribosomes. Indeed, given that TDP-43 target mRNAs represent a large proportion of cellular transcriptomes (Polimenidou et al., 2011), the lack of mRNA molecules may induce not only a worsening in protein aggregation (Maharana et al., 2018) but also the reorganization of ribosomes on available mRNAs. In line with this observation, we showed a widespread change in ribosomes recruitment on polysomes, which is also in agreement with the increase in detectable ribosomes in ALS models and in patients (Verheijen et al., 2014) and with the proposed trapping of ribosomes in stress granules (Anadolu et al., 2023). This change may then lead to an increased probability for non-target transcripts to be translated due to the increased ribosome availability, (Mills & Green 2017), which we observed using capacity monitor assays. Therefore, both global and specific protein levels may vary depending on available ribosomes, with the axons being particularly sensitive to changes in this balance. We supported this hypothesis at the mRNA-specific level for two transcripts (*Rrbp1*, *Vim*) that are transcriptionally dysregulated in post-mortem spinal cords of ALS patients (Krach et al., 2018).

In summary, our results provide a mechanistic insight into how TDP-43 proteinopathy disrupts both mRNA stability and translation, revealing a coordinated shift in cellular resource allocation. We show that TDP-43 target mRNAs undergo preferential destabilization and downregulation (Tank et al., 2018), accompanied by TDP-43 aggregation (Chen-Plotkin et al., 2010) and a compensatory increase in the translation of non-target mRNAs. This imbalance likely arises from enhanced ribosome availability following the loss of TDP-43 targets, leading to a global reorganization of ribosomes across the transcriptome. These results possibly reconciles the apparent inconsistencies in global, gene or cell compartment-specific protein levels already observed in ALS (Fiesel et al., 2012, Majumder et al., 2012; Majumder et al., 2016; Gao et al., 2021; Lehmkuhl et al., 2021; Russo et al., 2017; MacNair et al., 2016; Coyne et al., 2014; Altman et al., 2021; Chu et al., 2019; Higashi et al., 2013).

Together, these findings support a unifying model of ALS pathogenesis in which TDP-43-induced disruption of ribosome dynamics and mRNA homeostasis triggers a cascading imbalance in protein synthesis, offering a mechanistic framework with direct relevance to the early molecular events of the disease.

## MATERIAL AND METHODS

### Mouse model of ALS

The hTDP-43-Q331K mouse model of ALS (B6N.Cg-Tg (PrnpTARDBP*Q331K)103Dwc/J)) was purchased from The Jackson Laboratory and was generated on a C57BL6 background, as described in (Arnold et al., 2013). Non-transgenic C57BL/6N littermates were used as controls. Litters were genotyped using standard protocols. Mice were housed within the animal care facilities at the University of Trento under appropriate conditions. Animal breeding and procedures were conducted under appropriate project and personal license granted by the ethical committee of the University of Trento and were approved by the Italian Ministry of Health (D. Lgs no. 2014/26, implementation of the 2010/63/UE).

### Cultures of primary cortical neurons for microfluidic chambers

Primary cortical neurons were isolated from wild type C56BL/6J mice. The pregnant female was euthanized by CO2 at 14.5 days of gestation. The cortex was isolated from the embryos and purified from meninges, cut into small pieces and incubated for 5 minutes in DMEM/F12, 0.0125 M glucose, 1% P/S, 1 mM EDTA and then digested in trypsin by incubation in DMEM/F12, 0.0125 M glucose, 1% P/S, 0.25% trypsin, for 10 minutes at 37°C. After trypsin neutralisation in DMEM/F12, 0.0125 M glucose, 1% P/S, 20% FBS, samples were mechanically dissociated into single cells in DMEM/F12, 0.0125 M glucose, 1% P/S, 10% FBS, 4 KU/mL DNAseI, using fire-polished glass pipettes. The cells suspension was then filtered with a 70 μm cell strainer nylon filter, counted and centrifuged for 5 minutes at 163 g at RT to be resuspended in complete Neurobasal medium (Neurobasal, 2% B27 supplement, 18 mM HEPES, 1% P/S, 0.5 mM L-Glutamine). Cortical neurons were seeded on poly-D-lysine (PDL) and laminin coated glasses or plates at the desired concentration. Cortical neurons were maintained in culture at 37°C and 5% CO2 and half medium was changed every two days.

### Microfluidic chambers

Microfluidic chambers (SD450 Xona microfluidics) were used for cell body and axonal polysomal profiling and immunofluorescence. Glass coverslips were cleaned with 1 M HCI for 4 hours, extensively washed in milliQ water and dried in the oven. Next, the coverslips and the microfluidic chambers were sterilised in 70% ethanol and dried under a laminar flux hood at RT. The coverslips were coated overnight with 500 μg/mL poly-D-lysine and then assembled with the microfluidic device. The microfluidic chambers were equilibrated with PBS, followed by equilibration in the neurobasal medium. Cortical neurons resuspended in neurobasal medium at the concentration of 25,000 cells/μL were plated in a final amount of 250,000 cells/chamber following the SD450 Xona Microfluidics protocol. BDNF at a final concentration of 10 ng/mL was added every two days in the axonal compartment starting from DIV2.

### Production, titration and infection of lentiviral particles

Lentiviral particles were prepared as described previously (Amendola et al., 2005). Briefly HEK293T cells were transient co-transfected using the calcium-phosphate precipitation method with the transfer vectors, the MDLg/pRRE plasmid, the RSV-Rev plasmid and the MDLg plasmid encoding the G glycoprotein of the vesicular stomatitis virus. Cell supernatants containing lentiviral particles were collected 72 h after transfection, filtered and subjected to ultracentrifugation. The pellets were resuspended, divided into aliquots and stored at −80°C. To calculate the titer of lentiviral particles, cortical neurons were seeded in a 24-well plate and infected at 5 DIV with a serial dilution (from 10-3 to 10-7) of lentiviral particles overexpressing TRFP-TDP-43-WT, TRFP-TDP-43-A315T and TRFP. Uninfected cortical neurons were used as control. The experiment was repeated in triplicate for each condition. Cortical neurons were fixed at 9 DIV and processed for immunofluorescence with TRFP and TuJ1 primary antibodies. ArrayScan microscope (Thermo Fisher ArrayScan XTI HCA Reader) was used to acquire 20 random fields per well. For every field the TRFP-positive infected cells (red) and the total number of cells, stained with a nuclear marker (DAPI), were counted. Then, TRFP-positive cells and DAPI-positive cells coming from all fields were summed. A ratio between the number of the total TRFP-positive cells and the total DAPI-positive cells present in each well was calculated. To calculate the titer a percentage ratio between TRFP-positive cells and total DAPI positive cells between 0.1% and 10% (dynamic range) was chosen, and this formula was used: (Infected cells/Total cells) *Dilution factor*Number of seeded cells. In our experiments a MOI (multiplicity of infections) of 4 was used to ensure the infection of almost total cells without inducing cellular toxicity. Cortical neurons were infected at 5 DIV and treated, fixed or lysed at different DIV based on the experimental needs.

### Cell line cultures

Motor-neuron like cell line NSC-34 and hiPSC cell line were maintained in culture at 37°C with 5% of CO_2_. NSC-34 cells were cultured in Dulbecco’s Modified Eagle’s Medium (DMEM) with 4.5 g/l glucose supplemented with 2 mM glutamine, 10% FBS, 100 U/mL penicillin and 100 mg/mL streptomycin. hiPSC cells were seeded on Geltrex-coated 6-well plated and cultured in TeSR-E8 medium.

### Transduction and generation of NSC-34 inducible cell lines

The full-length human TDP-43 cDNA was amplified from SK-N-BE cells with AccuPrime DNA polymerase (Thermofisher Scientific) and cloned in SgfI and MluI sites of pCMV6-AN-His-HA plasmid (OriGene) to generate the vector pCMV6-His-HA-hTDP-43-WT, expressing the hTDP-43-WT gene with an N-terminal polyhistidine (His) and HA tag. The mutants hTDP-43-A315T, hTDP-43-Q331K, and hTDP-43-M337V were obtained via PCR-directed mutagenesis. For lentiviruses preparation, the His-HA tagged genes were excised from pCMV6-His-HA plasmids and subcloned into the BamHI and XhoI sites of the vector pENTR1A (Addgene). The resulting vectors were then recombined with pLenti CMV/TO Puro DEST (Addgene) to get the lentiviral vectors expressing His-HA-tagged hTDP-43-WT and its mutants under the control of a doxycycline-inducible promoter. Lentiviruses were then prepared in accordance with the protocols detailed by Campeau et al. 2009, meeting Biosafety Level 2 (BSL-2) requirements.

To generate inducible cell lines, NSC-34 cells were transduced with the pLentiCMV_TetR_Blast vector (Addgene), constitutively expressing the tetracycline (Tet) repressor under the control of a CMV promoter and selected for 7 days using 10 µg/ml Blasticidin (Sigma-Aldrich). The stable cells were infected with the lentiviral vectors expressing hTDP-43-WT and mutants in the presence of 4 µg/ml polybrene and selected by using 5 µg/ml puromycin.

### Subcellular fractionation

hiPSC were lysed with polysomal buffer containing 10 mM NaCl, 10 mM MgCl_2_, 10 mM Tris–HCl pH 7.5, 1% Triton X-100, 1% sodium deoxycholate, 0.005 U/µL DNAseI, 0.2 U/µL RNase inhibitor, 1 mM dithiothreitol and 10 μg/mL cycloheximide and protease-inhibitor EDTA-free. A few microlitres of lysate was retained for protein extraction (input). The sample was centrifuged for 90 min at 100,000 rpm using a TLA100.2 rotor in a Beckman Optima™ LE-80K Ultracentrifuge. The supernatant corresponds to the S100 fraction and does not contain ribosome-bound proteins. The pellet containing ribosomes (R pellet) was resuspended in high salt buffer containing 15 mM Tris-HCl pH 7.4, 500 mM KCl, 5 mM MgCl_2_, 2 mM dithiothreitol (DTT), 290 mM sucrose. Few microliters were maintained for protein extraction while the remaining volume was loaded on a discontinuous sucrose gradient (40% (w/V) sucrose in 15 mM Tris-HCl pH 7.4, 500 mM KCl, 5 mM MgCl_2_, 2 mM DTT (bottom layer); and 480 μl buffer 20% (w/V) sucrose, 15 mM Tris-HCl pH 7.4, 500 mM KCl, 5 mM MgCl_2_, 2 mM DTT (top layer) and ultracentrifuged at 4°C for 17 h using a TLS-5S rotor at 33,000 rpm in Beckman Optima™ LE-80K ultracentrifuge. The supernatant (WS) contained proteins loosely associated with ribosomes and the pellet contained washed ribosomes (WR). Pellet was dissolved in sample buffer and proteins in the other fractions extracted using methanol-chloroform. For RNAse treatment, cytoplasmic lysate was treated with RNAse I (10U per unit absorbance at 260 nm of lysate) for 45 minutes at RT, before applying the same protocol. Western blot analysis was done using the primary antibodies detailed in **Supplementary Table 1.**

### Ribosome profiling and active ribosome profiling

Cytoplasmic lysates from control and ALS mouse primary cortical neurons were prepared as previously described (Bernabò et al. 2017). The RNA content of each lysate was assessed by measuring the absorbance at 260 nm with the Thermo Scientific NanoDrop. The NaCl concentration was adjusted to 100 mM. If not processed immediately, samples were stored at −80 °C. Lysates were treated with RNase I (2/3 Invitrogen and 1/3 BioResearch Technologies) using 7.5 U / a.u. Abs260 nm of the lysate and incubated at room temperature for 45 minutes. The reaction was stopped by adding 100 U of SUPERase-In RNase inhibitor (Thermo Fisher Scientific). Polysome profiling was performed as previously described (Lauria et al. 2020). Fractions corresponding to the 80S monosome were collected. For RiboSeq, the hyper-resolution RiboLace 360 Gel Free Kit (Immagina BioTechnology, catalog n. 360SQ-12) was used for the library preparation according to the manufacturer’s instructions. For active RiboSeq, the RiboLace kit (Immagina BioTechnology) was used for ribosome purification and library preparation according to the manufacturer’s instructions. Eventually, RiboSeq and active RiboSeq libraries were quantified using High Sensitivity DNA kit (Agilent Technologies) on the Agilent 2100 Bioanalyzer. Single read sequencing 100 cycles was performed with the Illumina NovaSeq 6000 by the Next Generation Sequencing Facility (CIBIO, University of Trento, Italy).

### Miniaturised polysome profiling from cortical neurons cultured in microfluidic chambers

The compartmentalised axons and cell bodies were lysed directly in the corresponding compartments of the microfluidic chamber by fluxing several times 100 μL of polysome lysis buffer (10 mM NaCl, 10 mM MgCl_2_, 10 mM Tris–HCl pH 7.5, 1% Triton X-100, 1% sodium deoxycholate, 0.005 U/mL DNAseI, 0.2 U/µL RNase inhibitor, 1 mM DTT and 10 μg/mL cycloheximide). The lysates were centrifuged at 14000 g for 5 minutes at 4°C to pellet membrane debris, nuclei and mitochondria. The cell body supernatant was incubated 20 minutes before storage at −80°C. For polysome profiling the lysates were loaded onto a 15–50% linear sucrose gradient formed in 2.2 mL polyallomer tubes. Sucrose buffers were prepared in 100 mM NaCl, 10 mM MgCl_2_, 30 mM Tris–HCl, pH 7.5. Sucrose gradients were centrifuged in a Beckman Optima™ TL Ultracentrifuge for 90 minutes at 180,000 g at 4°C using a Beckman TLS-55 swinging rotor. To collect sucrose fractions and measure the absorbance at 254 nm, a Teledyne Isco model 160 gradient analyser equipped with a UA-6 UV/VIS detector was employed. The resulting 200 μl fractions were directly used to extract RNA samples or stored at −80°C.

### RNA purification from polysomal profiles, library preparation

For studying both the transcriptional and the translational levels of the axons and cells, total cytosolic RNA (pool of all the fractions) and polysomal RNA (collection of fractions corresponding to the polysomal peaks) were purified from the polysomal profiles. Both total and polysomal RNA extractions were performed with Direct-zol^™^ RNA purification Kit (Zymo Research). For RNPs/granule transcriptome and translatome analysis at subcellular resolution, after sucrose gradient fractionation, sucrose fractions not associated to polysomes containing RNPs, ribosomal subunits and 80S were pooled in the RNPs/granule fraction. Sucrose fractions containing polysomes were pooled into the “polysome fraction”. Trizol was used to extract the RNA. After purification, the RNA was resuspended in DNAse-RNAse nuclease free-water and quantified at the Agilent 2100 Bionalyzer by using the Agilent RNA 6000 Pico kit. The libraries were obtained using Ovation® Single Cell RNA-Seq System (Nugen) or Ovation® SoLo® RNA-Seq library preparation kit (Tecan) according to the manufacturer’s protocol. The qualitative and quantitative control of the libraries was performed with High Sensitivity DNA Chip (Agilent Technologies) before sequencing. Single read sequencing 100 cycles was performed with the Illumina NovaSeq 6000 by the Next Generation Sequencing Facility (CIBIO, University of Trento, Italy).

### RT-PCR from RNP/granule and polysomal RNA in axons and cell bodies

For PCR, reverse transcription reaction was performed on 100 pg of RNP/granule and polysomal RNA of cell body and axonal compartments, using the RevertAid First Strand cDNA synthesis kit (Thermo Fisher Scientific). The PCR reaction was performed on 1 μL cDNA for each sample, using Taq DNA Polymerase (Invitrogen), 10X PCR Buffer – Mg, 50 mM MgCl_2_, 10 mM dNTPs and 10 μM FW and RV primers of the selected genes. PCR was performed using C1000 Touch Thermal Cycler (Biorad). The PCR program was set as follows: initial denaturation step at 94°C for 3 minutes, 35-40 cycles composed of a denaturation step at 94°C for 45 seconds, annealing step at 58-60°C for 30 seconds and extension step at 72°C for 90 second. Final extension was performed at 72°C for 10 minutes. The PCR reactions were run on 8% polyacrylamide gel in TAE and amplicons visualized at the ChemiDoc MP Imaging System (Biorad). Primers are listed in **Supplementary Table 2**.

### Co-sedimentation profiles of proteins from cell bodies and axonal polysome profiling

After polysome profiling, the proteins were extracted fraction by fraction with methanol-chloroform method (Bernabò et al., 2017). After resuspension in milliQ water, 1 μL of each fraction was spotted onto a nitrocellulose membrane for dot blot analysis. Membranes were left drying for 30-45 min at room temperature and incubated in a blocking solution for 4 hours at RT in rotation. Next, membranes were incubated with primary antibody overnight at 4°C, under orbital agitation. The following day, membranes were washed twice in TBS-Tween for 30 minutes and incubated at RT with secondary antibody for 1h and 30 minutes. After antibody removal, two washes with TBS-Tween were followed by a final wash with TBS (30 minutes/wash). The reaction was developed by using SuperSignal™ West Femto Maximum Sensitivity Substrate (ThermoScientific) and the images acquired with ChemiDoc MP Imaging System (Biorad). To obtain the relative distribution of proteins along the profile, the percentage of each protein fraction by fraction was obtained as follows:

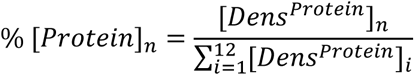

where *n* is the number of a specific fraction between 1 and 12 and % [*Protein*]_*n*_ is the percentage of the protein of interest in the fraction of interest, and *Dens*^*Protein*^ is the densitometry of the protein signal.

### Polysome profiling from cell lines and tissues and protein extraction

Doxycycline-inducible NSC-34 cells were seeded at 1.5 × 10^6^ cells/10 cm dish and treated with 2 µg/ml doxycycline (Clontech) for 48h to express His-HA-tagged hTDP-43 WT or mutants (hTDP-43-A315T, hTDP-43-Q331K, and hTDP-43-M337V). After reaching 80-90% of confluence, cells were treated with 10 μg/mL cycloheximide for 3 minutes at 37°C. Cytoplasmic lysates were obtained as described earlier (Bernabò et al., 2017). The lysate was then centrifuged at 20000 g for 5 min at 4°C to allow the sedimentation of membrane debris, nuclei and mitochondria. The supernatant was recovered and after 20 min on ice was stored at −80°C or directly transferred onto a 10–40% linear sucrose gradient as described in (Lauria et al., 2020) and centrifuged in a Beckman Optima™ LE-80K Ultracentrifuge for 90 min at 180,000 g at 4°C using a swinging rotor. To collect sucrose fractions and measure the absorbance at 254 nm, a Teledyne Isco model 160 gradient analyser equipped with a UA-6 UV/VIS detector was employed. Each fraction was stored at - 80°C for protein extraction. Mouse cortex and spinal cords from the hTDP-43-Q331K mouse model of ALS and controls were dissected and pulverised in a mortar with a pestle under liquid nitrogen and the powder was lysed as described in (Lauria et al., 2020). Tissue debris were removed by centrifugation at 18.000 g for 1 min at 4°C. Nuclei were removed from the pre-cleared lysate by a second centrifugation at 13.200 g for 5 min at 4°C and incubated on ice for 30 min. The polysome-containing lysate was stored at −80°C or directly layered onto a linear sucrose gradient as previously described. Protein isolation from both cell cultures and tissues was performed fraction by fraction using classical methanol/chloroform precipitation. The protein pellet was resuspended in 60 μl Electrophoresis Sample Buffer (Santa Cruz) before immunoblotting.

### Fraction of ribosomes in polysomes (FRP)

From the chromatogram of the polysome profiles the fraction of ribosomes in polysomes (FRP) was obtained as follows:

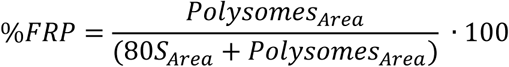

where *Polysomes*_*Area*_ is the area under the peaks of polysomes, the 80*S*_*Area*_ is the area under the 80S. Statistical analysis of data was carried out by using Student’s T-test for identification of difference between two groups.

### Total protein purification and Western blotting from cortex and spinal cords of the mouse model of ALS

Frozen spinal cord and cortex tissues from control or ALS mice were pulverised using mortar and pestle working under liquid nitrogen during the whole procedure. The powder was lysed in RIPA buffer (50 mM Tris-HCl pH 8.0, 150 mM NaCl, 1% NP40, 0.5% sodium-deoxycholate, 2% SDS) supplemented with protease inhibitor and homogenised with a Kimble® 749540-0000 Pellet Pestle® Cordless Motor. The samples were incubated on ice for 15 min and centrifuged at 15,000 rpm in F-45-30-11 rotor at 4°C for 15 min to pellet cell debris. The supernatants were collected and stored at −80C. Protein quantification was performed by using the DC assay kit (BioRad). For total protein analysis, the same quantity of proteins (20–30μg) for each sample was used. Protein samples were diluted in 2X Electrophoresis Sample Buffer (Santa Cruz) and milliQ water to reach a final loading volume of 20 μL and loaded on polyacrylamide gels (8%, 10% or 12%). After total protein evaluation by ponceau staining, WB analysis was performed as previously described. The list of primary and secondary antibodies used is in **Supplementary Table 1**. To determine the Fold Change of protein expression between ALS and control conditions, the signals’ intensities of the Western Blot bands were measured using the ImageJ software (https://imagej.nih.gov/). The signal’s intensity of each protein in each replicate was normalised to the respective intensity of ActB protein (housekeeping). Thus, the Normalized Protein Expression was defined as follows:

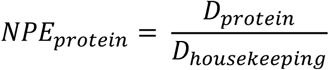

where *D*_*protein*_ is the signal’s intensity relative to the protein in a replicate, *D*_*housekeeping*_ is the signal’s intensity of the Western Blot band corresponding to ActB in the replicate. For each protein, the mean of the NPE obtained from the three control replicates was calculated as follows:

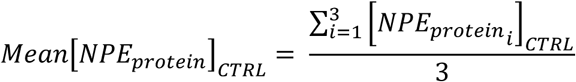

Lastly, the mean NPE control, calculated for a protein, was used to compute the FC of protein expression in ALS with respect to the control condition:

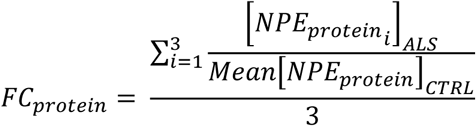

Statistical analysis of data was carried out by using Student’s T-test for identification of difference in total protein expression between two groups.

### RT-qPCR from total RNA and co-sedimentation analysis of mRNAs

Polysomal profiling from frozen spinal cord and cortex tissues, derived from (3-months old) ALS (TDP-43-Q331K) and control mice, was performed as described previously (Bernabò, et al. 2017). Total cytosolic RNA (pool of all the sucrose fractions) and RNA from each sucrose fraction were extracted using phenol-chloroform as described previously (Tebaldi et al., 2018). Before the reverse transcription reaction, all samples were treated with DNAse. Equal volumes of RNA from sucrose fractions were used for cDNA synthesis using the RevertAid First Strand cDNA synthesis kit (Thermo Fisher Scientific). For total cytoplasmic RNA 500 ng were used for cDNA synthesis using the RevertAid First Strand cDNA synthesis kit (Thermo Fisher Scientific). In both cases, cDNA synthesis reaction was performed by using OligodT. qPCR was performed using the CFX Connect Real-Time PCR Detection System (BioRad) using the qPCRBIO SyGreen Mix Separate-ROX (PCR Biosystem). The list of primers is in **Supplementary Table 2**. For qPCR data analysis to obtain the co-sedimentation profiles of the mRNAs of interest, the percentage of each transcript’s distribution along the profile was obtained using the following formula in the case of qPCR:

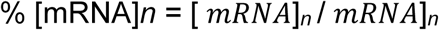

where n is the number of the fraction and %[mRNA]n is the percentage of mRNA of choice in each fraction, C_t_ mRNA is the Ct for each RNA in each fraction. For the differential expression analysis of the mRNAs of interest, the qPCR results of gene expression were normalised to ActB expression level. Precisely, the ΔCT for each transcript was calculated as CTgene – CTActB. Then, the log2FC was calculated from ΔΔCT values obtained comparing the expression of the gene of interest between TDP-43-WT (Control) or TDP-43-Q331K (ALS) condition.

### Ribosome run-off experiments and calculation of ribosome kinetics

Primary cortical neurons were extracted from mouse embryos (E15.5) and maintained in culture, as described earlier. At DIV 7, cells seeded in 10 cm Petri dishes (5 million cells/dish) were treated with Harringtonine (Abcam) in Neurobasal medium (GibcoTM) at a final concentration of 2 μg/mL. After harringtonine addition, the dishes were gently mixed and were incubated at 37°C for 0, 1.5, 3 and 10 min. Sharply, at the end of the harringtonine-incubation time, cycloheximide was added to the cell culture medium to a final concentration of 10 μg/mL to stabilise elongating ribosomes. Finally, a cytoplasmic lysate was obtained as described earlier and analysed by polysomal profiling. For each time point of harringtonine treatment (0, 1.5, 3, 10 min), the mean FRP value among the replicates was calculated. The obtained mean FRP values were fitted with the following exponential decay function, separately for control and ALS conditions:

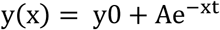

The ribosome kinetics was measured as the reciprocal of the decay constant (τ = 1/t), thus expressed as minutes^-1^ (min^-1^).

### Puromycilation in PCNs cultured in microfluidic chambers, imaging and particle analysis

The puromycilation reaction was performed according to (David et al., 2012). Mouse cortical neurons (DIV10) were incubated with 100 μg/ml cycloheximide for 15 minutes at 37°C. Next, 100 μg/ml puromycin was added and kept for 5 minutes at 37°C. Cells were washed on ice with cold HBS (119 mM NaCl, 5 mM KCl, 2 mM CaCl_2_, 2 mM MgCl_2_, 30 mM glucose, cycloheximide 100 μg/ml and 0.0003% digitonin, 25 mM Hepes, pH 7.3) for 2 min. A second wash was performed with the same buffer without digitonin to eliminate the detergent. Cells were then fixed 15 minutes with PBS solution containing 4% paraformaldehyde, 4% sucrose and 100 μg/ml cycloheximide. After 3 washes with PBS, cells were permeabilized and blocked with 5% BSA, 0.1% Triton X-100 in PBS for 30 minutes and then incubated with primary antibodies (**Supplementary Table 1**) for 1 h. After 3 washes with PBS, the coverslips were incubated with a convenient secondary antibody for 1 hour. Cells were then washed with PBS and microfluidic chambers detached from the coverslip to be mounted with Prolong Diamond Antifade Mountant containing DAPI (Invitrogen). Images were acquired using a Leica TCS SP8 microscope. 4096 x 4096 pixel (pixel size/ voxel size: 0.284 μm) images of whole chambers were acquired using HC PL FLUORAR 10x/0.30 dry objective. 2048 x 2048 pixels (pixel size/ voxel size: 0.09 μm) images of cell bodies and axons were acquired using HC PL APO CS2 63x/1.4 NA oil objective. Samples were illuminated using Diode 405, Argon and DPSS 561 lasers, for excitation at 405 nm, 488 nm and 561 nm wavelengths, respectively. Images were acquired with PMTs detectors using the following parameters: Alexa Fluor 488: 500–540 nm, TRFP: 566-650 nm, DAPI: 410-475 nm emission wavelength ranges. Z-stack imaging was performed on cell bodies, axons in the microgrooves and distal axons. Images of Vim and Rrbp1 immunofluorescence were taken on a Leica TCS SP8 microscope with the same parameters. For Gapdh, samples were imaged on a Leica SP5 microscope. Images at 4096 x 4096 pixels (pixel size/ voxel size: 0.378 μm) were acquired from the whole chambers using HC PL FLUORAR 10x/0.30 dry objective. Images at 2048 x 2048 pixels (pixel size/ voxel size: 0.12 μm) for the cell bodies and axons in the microgrooves were acquired using HCX PL APO CS 63x/1.4 NA oil objective. Samples were excited using Argon and HeNe lasers and excitation wavelengths were set at 488 nm and 543 nm, respectively. Images were acquired with PMTs detectors using the following parameters: Alexa Fluor 488 500–540 nm and TRFP 566-650 nm. Z-stack imaging was performed in cell bodies and axons in the microgrooves. The images were analyzed with ImageJ-software (https://imagej.nih.gov/ij/). To count puromycin positive foci and measure their area and perimeter, images of axon in the microgrooves were processed to subtract the background and a threshold was set at 1% before producing a binary image. Regions of interests (ROI) for axons were captured at 112 x 2048 pixels (10 μm x 185 μm). For particle analysis a lower size threshold for particles count was set to 20 pixel units. Area and perimeter were measured for all the included particles. Particles with area < 0.3 μm^2^ were excluded from the analysis.

### Fluorescence intensity analysis of cortical neurons in microfluidic chambers

Puromycin, Gapdh, Vim and Rrbp1 fluorescence intensities were analysed with the ImageJ-software (https://imagej.nih.gov/ij/). Stacks of each image were Z-projected with a convenient plugin and fluorescence measured as integrated density. In the images of the cell body the fluorescence was measured relatively to the number of cells in each ROI. ROI fluorescence was normalised to the background intensity and the results expressed as variations with respect to the TRFP control condition that was set to 1.

### Translational burden analysis

NSC-34 cells were maintained in Dulbecco’s modified Eagle medium (DMEM, Gibco) supplemented with 10% FBS (Atlanta BIO), 1% penicillin/streptomycin/L-glutamine (Sigma-Aldrich), and 1% non-essential amino acids (HyClone) at 37 °C and 5% CO_2_. For transfection, 3.75*10^4^ cells per well were transfected in 48-well plates. The transfection solution was prepared using polyethylenimine (PEI) “MAX” (Mw 40,000, Polysciences, Inc.) in a 1:2 (μg DNA to μg PEI) ratio with a total of 150 ng of plasmid DNA per well. Both DNA and PEI were diluted in 12.5 μL of Opti-MEM I reduced serum media (Gibco) per well before being mixed and incubated for 25 min prior to addition to the cells. To determine the translational burden, cells were analysed with a BD Celesta™ cell analyzer (BD Biosciences) using 488 and 561 lasers. For each sample >30,000 events were collected and fluorescence data were acquired with the following cytometer settings: 405 nm laser and 450/40 bandpass filter for EBFP, 488 nm laser and 530/30 nm bandpass filter for EGFP, 561 nm laser and 610/20 nm filter for TRFP. Transfected cells were washed with DPBS, detached with 15 μL of Trypsin-EDTA (0.25%), and resuspended in 125 μL of DPBS + 1% FBS + 0.2 mM EDTA (Thermo Fisher). Fluorescence intensity in arbitrary units (a.u.) was used as a measure of protein expression. For each experiment a compensation matrix was created using unstained (wild type cells), and single-color controls (EBFP only, TRFP only, EGFP only). Live cell population and single cells were selected according to FCS/SSC parameters. Data analysis was performed with FlowJo V.10.

### Data analysis

#### Preprocessing and alignment of ribosome profiling data

RiboSeq and active RiboSeq data were processed following the instructions provided by Immagina BioTechnology. Briefly, after adapter removal (TCTCCTTGCATAATCACCAACC), the UMIs (first 4 and last 4 nucleotides) and then the barcodes (first 8 nucleotides) were extracted, and only reads starting with a T, which was trimmed, and longer than 15 nucleotides were retained (CutAdapt v3.7; UMI-Tools v1.1.2). Reads mapping on the collection of M. musculus rRNAs (from the SILVA rRNA database) and tRNAs (from the Genomic tRNA database) were discarded and the remaining reads were mapped on the reference genome GRCm39 with STAR (v2.7.10a), using the Gencode vM32 gene annotation.

#### Ribosome profiling analyses

Positional analyses were performed using the riboWaltz R package v2.0 (Lauria et al., 2018). Differential ribosome occupancy analyses are based on the number of P-sites falling in the coding sequence of each transcript, as computed by riboWaltz.

#### Preprocessing and alignment of RNA-Seq data

Reads were processed by removing the 3’ Illumina Universal Adapter (AGATCGGAAGAG), trimming the first 8 nucleotides and discarding reads shorter than 40 nucleotides (Cutadapt v2.5). Processed reads were aligned to the mouse genome (GRCm38.p6) with STAR (v2.5.3a), using the Gencode M17 gene annotation, based on ENSEMBL release 92. A maximum of 10 multiple alignments for each read was allowed. Reads mapping at the same positions, but the one with the highest mapping quality score, were removed with Picard Tools MarkDuplicates (v2.18.9, REMOVE_DUPLICATES=true). Counts of mapped reads per gene were retrieved by HTSeq (v0.11.0, --stranded=no). All programs were used with default settings unless otherwise specified.

#### Differential analyses

To remove size or compositional differences between libraries, gene expression levels were normalised among replicates using the trimmed mean of M-values normalisation method (TMM) implemented in the edgeR Bioconductor package. Only genes with FPKM ⩾ 10 in all replicates of at least 1 sample were kept for subsequent analysis. Differentially expressed genes, cellular and axonal enriched genes and genes with decreased and increased TE were detected with edgeR (glmQLFTest function) with a triple threshold: i) absolute value of log2 fold change, log2 fold enrichment and fold translational efficiency ⩾ 0.75; ii) p-value representing the statistical significance ⩽ 0.05; iii) average CPM among replicates of each sample ⩾ 0.05.

#### Network analyses

Protein-protein interactions were downloaded from the STRING database (v 11.0, Mus Musculus dataset). Only interactions with interaction score ⩾ 0.3 were considered (this threshold includes medium and high confidence interactions). Network analysis was performed with the igraph R package and protein communities within the network were identified with the cluster_fast_greedy function.

#### Functional enrichment analyses

Annotation enrichment analysis with Gene Ontology terms, REACTOME and KEGG pathways were performed with the clusterProfiler Bioconductor package.

#### Analysis of TDP-43 CLIP data

CLIP-Seq data retrieved from Polymenidou et al., 2011 were aligned to the mouse transcriptome using the Gencode M17 annotation, based on ENSEMBL release 92.

#### Definition of TDP-43 targets

TDP-43 targets were defined according to the presence of TDP-43 binding motifs, retrieved from Polymenidou et al., 2011, on the 3’ UTR of annotated protein codings. Genes not including any motifs were labelled as non-target RNAs.

#### Enrichment analyses

Enrichment of TDP-43 target genes defined in this work and destabilised genes (retrieved from Tank et al., 2018) among DEGs identified upon TDP-43-WT or TDP-43-A315T expression were performed by Fisher’s exact test. The set of genes with FPKM ⩾ 10 in all replicates of at least 1 sample were used as background.

## Data availability

RNA-Seq data generated by the current study have been deposited in the Gene Expression Omnibus (GEO) under the accession code GSE239419. All other data supporting the findings of this study are available from the corresponding author on reasonable request. Source data are provided with this paper.

## Supplementary Data

Supplementary Data are available at NAR Online

## Acknowledgements

We thank the staff at the Next Generation Sequencing Facility (NGS) at Department CIBIO University of Trento for technical support.

## Funding

This work was supported by Provincia Autonoma di Trento, Italy (AxonomiX research project granted to A.Q. and G.V.), the ALS Association U.S.A. (pilot project 2018; granted to G.V.), the ARISLA Foundation (AxRibALS, call 2017 granted to G.G.C. and G.V.). We also acknowledge financial support from IMMAGINA Biotechnology (Italy).

## Author contributions

G.V. conceived and designed the study and directed the research. F.L., G.T., T.T. and E.B. processed all sequencing data and performed the analyses; F.M., M.M., I.B., M.S., C.P., L.L., R.A., A.B. and M.d’A. performed experimental work. A.P., L.C. and G.G.C. optimised and produced the viral particles for the expression of TDP-43-WT and TDP-43-A315T. D.P. and A.P. produced the cellular lysates from hiPSCs. D.P. and A.Q. developed the NSC-34 stable and inducible cell lines. L.D. and M.B. dissected all tissues from the mouse model. F.C. and V.S. produced plasmids and performed the translational burden experiments. A.Q, G.G.C. and G.V. obtained the funding. G.V., F.M., F.L., C.P. and R.A. wrote the manuscript, with input from all authors.

## Declaration of interests

G.V. has been a scientific advisor to IMMAGINA Biotechnology until November 2022. M.C. is the CEO of IMMAGINA Biotechnology. The other authors declare no competing interests.

## SUPPLEMENTARY FIGURES

**Supplementary Figure 1.**
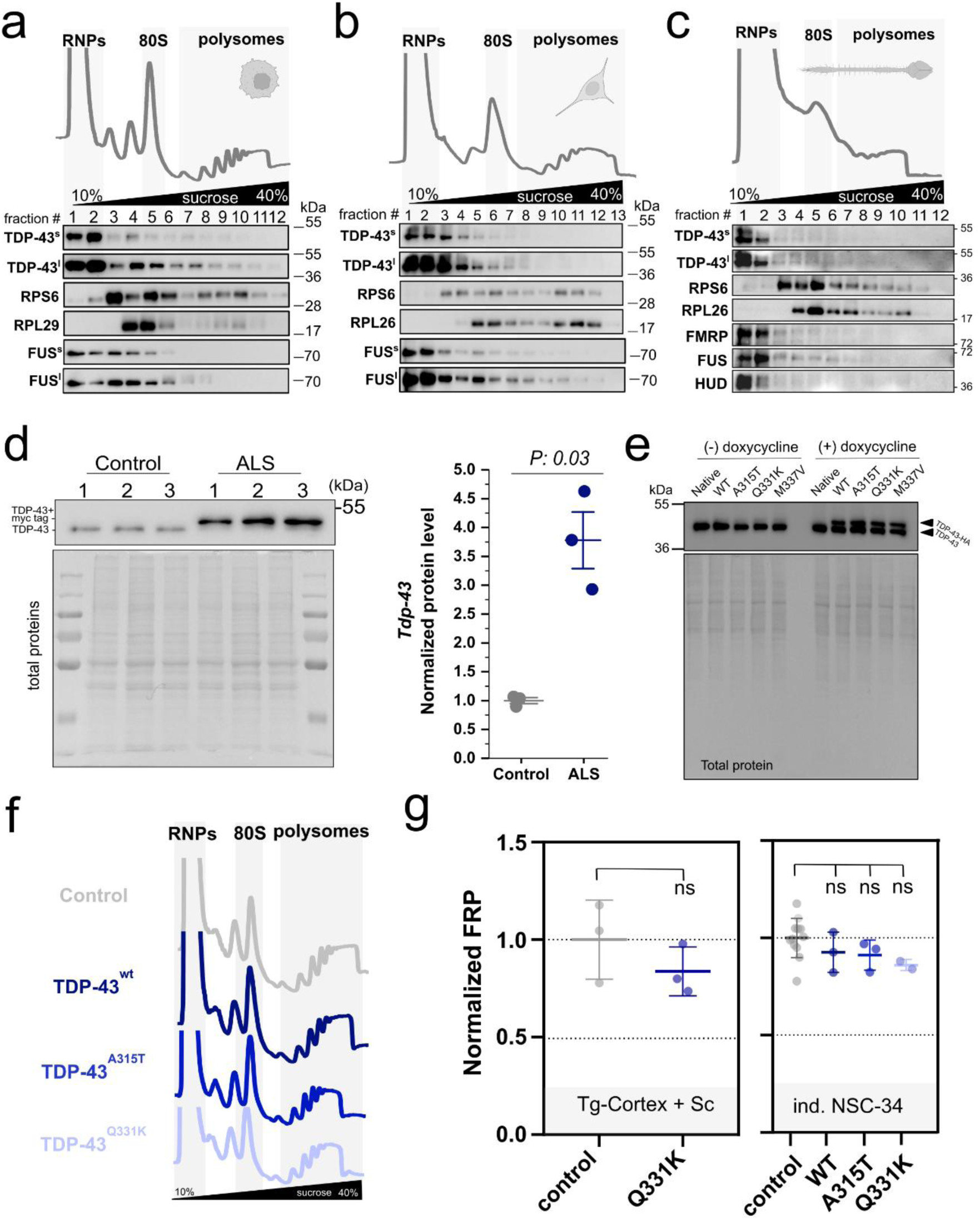
(**a-c**) Representative polysome profiles (upper panels) and representative co-sedimentation profiles (lower panels) of TDP-43 (s: short exposure, l: long exposure), ribosome markers (RPS6, RPL29 or RPL26), and RNA-binding proteins (FUS, FMRP, and HUD) performed in (a) hiPSCs, (b) NSC-34 cell lines and (c) murine central nervous system (CNS). hIPSC, NSC34 and mouse cortex+SC illustrations created in BioRender. (**d**) Left panel: western blot analysis of TDP-43 and hTDP-43-Q331K in the cortex and spinal cord from 3 months-old non-transgenic controls and hTDP-43-Q331K mouse model of ALS. Total proteins were assessed using ponceau dyeing of the membrane. Lanes 1-3 represent biologically independent replicates. Right panel: quantification of TDP-43 protein levels in 3 months-old control and hTDP-43-Q331K mouse model of ALS after normalisation for total protein abundance. Data are mean ± SEM of n=3 biologically independent replicates. Differences were assessed with two-tailed T-test. P-value is reported in figure (**e**) Western blot analysis of transgene (human) and endogenous (murine) TDP-43 in NCS-34 cells stably transduced to express His-HA-tagged hTDP-43-WT and mutants A315T, Q331K and M337V after induction with doxycycline. (**f**) Examples of polysome profiles from NSC-34 expressing His-HA-tagged hTDP-43-WT or two hTDP-43 mutants (A315T and Q331K). (**g**) Fraction of ribosomes in polysomes (FRP) in the cortex and spinal cord of control and transgenic mice expressing hTDP-43-Q331K (left panel) and doxycycline-inducible NSC-34 cell lines expressing His-HA-tagged hTDP-43-WT, hTDP-43-A315T, and hTDP-43-Q331K (right panel). Data are mean ± SEM of n=3-9 biologically independent replicates. No significant changes according to two-tailed T-test were detected.

**Supplementary Figure 2.**
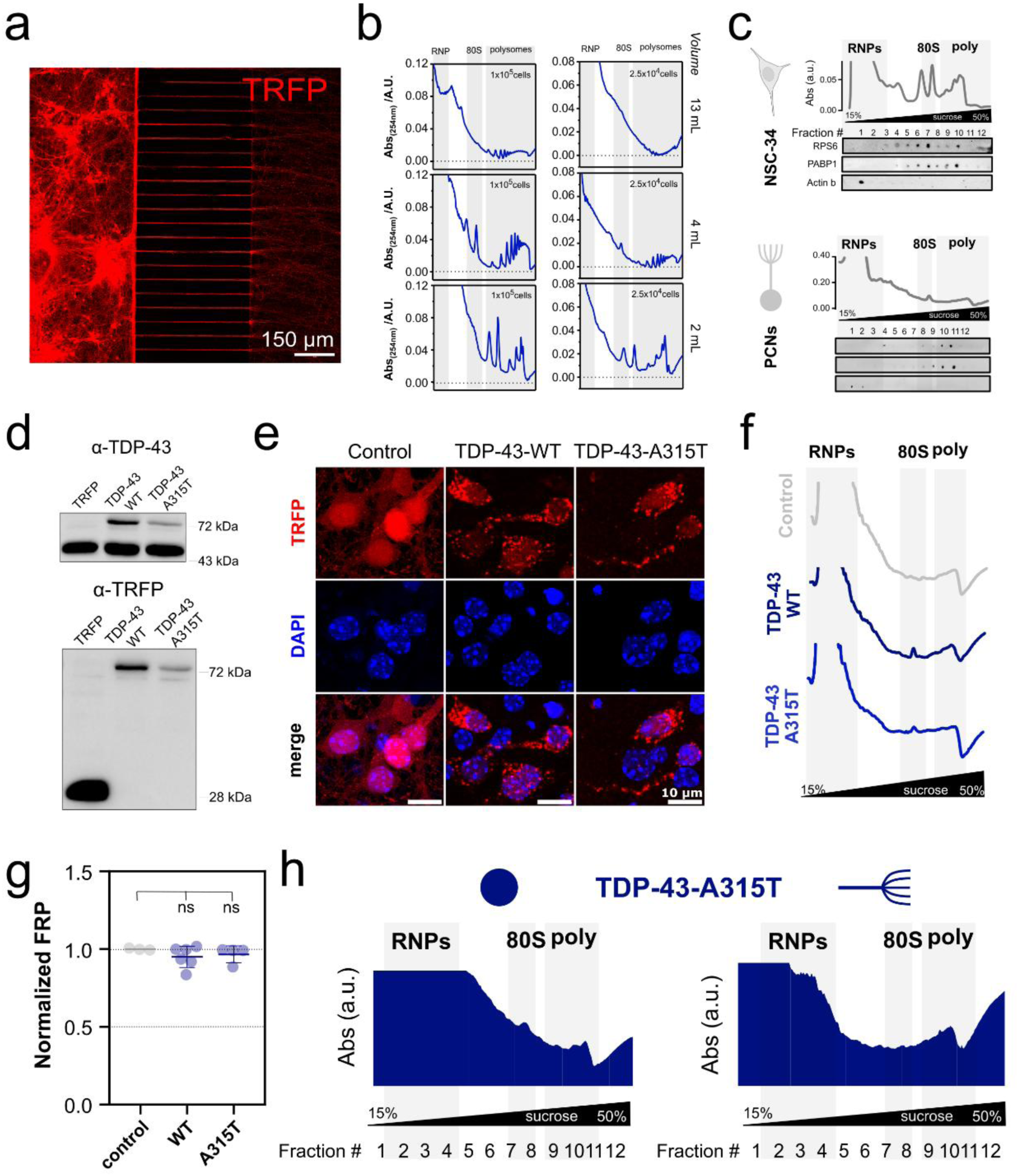
(**a**) Representative immunofluorescence image of mouse PCNs expressing TRFP cultured in microfluidic chambers to compartmentalise cell bodies (left) and axons (right). (**b**) Polysomal purification optimization in 15%-50% sucrose gradients formed in different tubes and with a decreasing number of MCF7 cells. The miniaturised 2 mL sucrose gradient efficiently allows the sedimentation and isolation of polysomes. (**c**) representative polysomal profile and dot blot co-sedimentation profiles of ribosomal markers obtained with miniaturised polysome profiling in control NSC-34 (left panels) and in wild type PCNs (right panels). PABP was used as a control for RNA-dependent interactions and Actin B was used as negative control. NSC34 illustrations created in BioRender. (**d**) Western blot of total protein lysates obtained from murine PCNs infected with TRFP, TRFP-TDP-43-WT and TRFP-TDP-43-A315T to monitor the expression levels of endogenous TDP-43 and exogenous TRFP-TDP-43-WT and TRFP-TDP-43-A315T proteins. (**e**) Immunofluorescence of mouse PCNs expressing TRFP-TDP-43-WT, TRFP-TDP-43-A315T, and only TRFP as control. Antibodies against TRFP were used to evaluate TDP-43-positive inclusions in the cytoplasm, compared to nuclear marker (DAPI). (**f**) Examples of polysome profiles from PCNs infected with TRFP as control, and PCNs expressing TRFP-TDP-43-WT and TRFP-TDP-43-A315T. (**g**) Fraction of ribosomes in polysomes (FRP) in PCNs expressing TRFP-TDP-43-WT, TRFP-TDP-43-A315T and TRPF as control. Data are mean ± SEM of n=3-6 biologically independent replicates. No significant changes according to two-tailed T-test were detected. (**h**) Representative profiles obtained with miniaturised polysome profiling in axons and cell body of murine PCNs expressing TRFP-TDP-43-A315T.

**Supplementary Figure 3.**
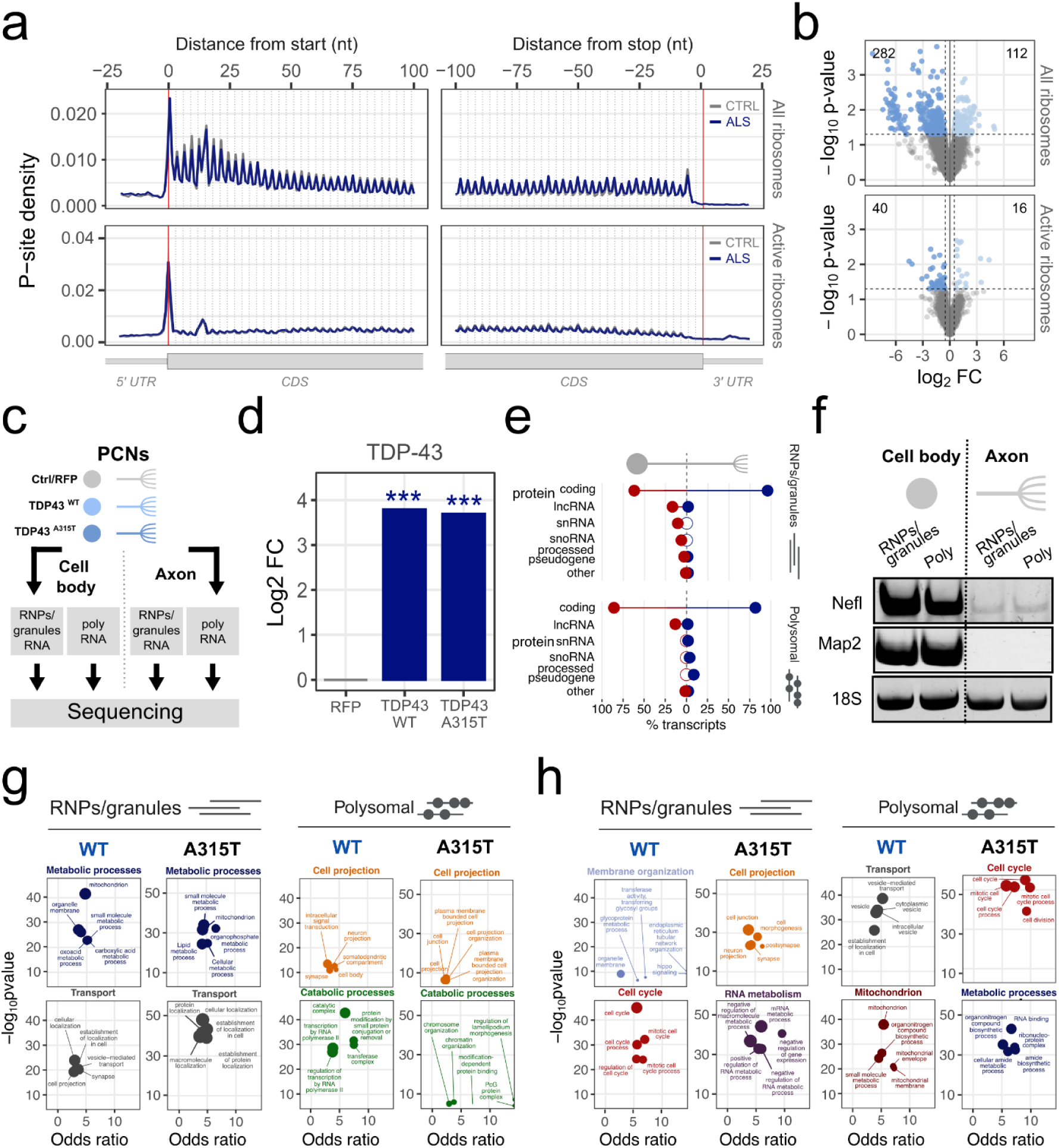
(**a**) Overlay meta-profiles based on the localization of active and inactive ribosomes (top) and only active ribosomes (bottom) in ctrl (gray) and ALS (blue). Data are mean ± SEM of n=3 biologically independent replicates. (**b**) Volcano plot displaying variations in the ribosome occupancy according to active and inactive ribosomes (upper panel) and only active ribosomes (bottom panel). mRNAs displaying significant variations are coloured according to the direction of the change: decreased ribosome occupancy (blue) and increased ribosome occupancy (light blue). The number of transcripts is reported above. (**c**) Experimental design for the isolation of RNAs not associated with polysomes (RNPs/granules) and associated with polysomes (polysomal RNA) in cell bodies and axons. from PCNs expressing TDP-43-WT, TDP-43-A315T, and TRFP as a control, cultured in microfluidic chambers. (**d**) Level of cellular polysomal TDP-43 RNA in the cell bodies of neurons expressing TDP-43-WT and TDP-43-A315T with respect to TRFP. Differences were assessed with two-tailed T-test (P-value: *** p <0.001). (**e**) Percentage of biotypes of free (top panel) and polysomal (bottom panels) RNAs found to be enriched in the cell body (red) or in axonal (blue) compartment in control cortical neurons. (**f**) RT-PCR of free and polysomal RNA for the neuronal cytoplasmatic Nfl, the dendrite marker Map2, and the ribosome marker 18S in control PCNs expressing TRFP. (**g**) Functional enrichment analysis of the top two communities based on the interaction network of proteins encoded by RNPs/granules (left panel) and polysomal (right panel) genes enriched in the axon, shared across PCNs upon TDP-43-WT and TDP-43-A315T expression. (**h**) Functional enrichment analysis of the top two communities based on the interaction network of proteins encoded by RNPs/granules (left panel) and polysomal (right panel) genes enriched in the axon, not shared across PCNs upon TDP-43-WT and TDP-43-A315T expression.

**Supplementary Fig. 4.**
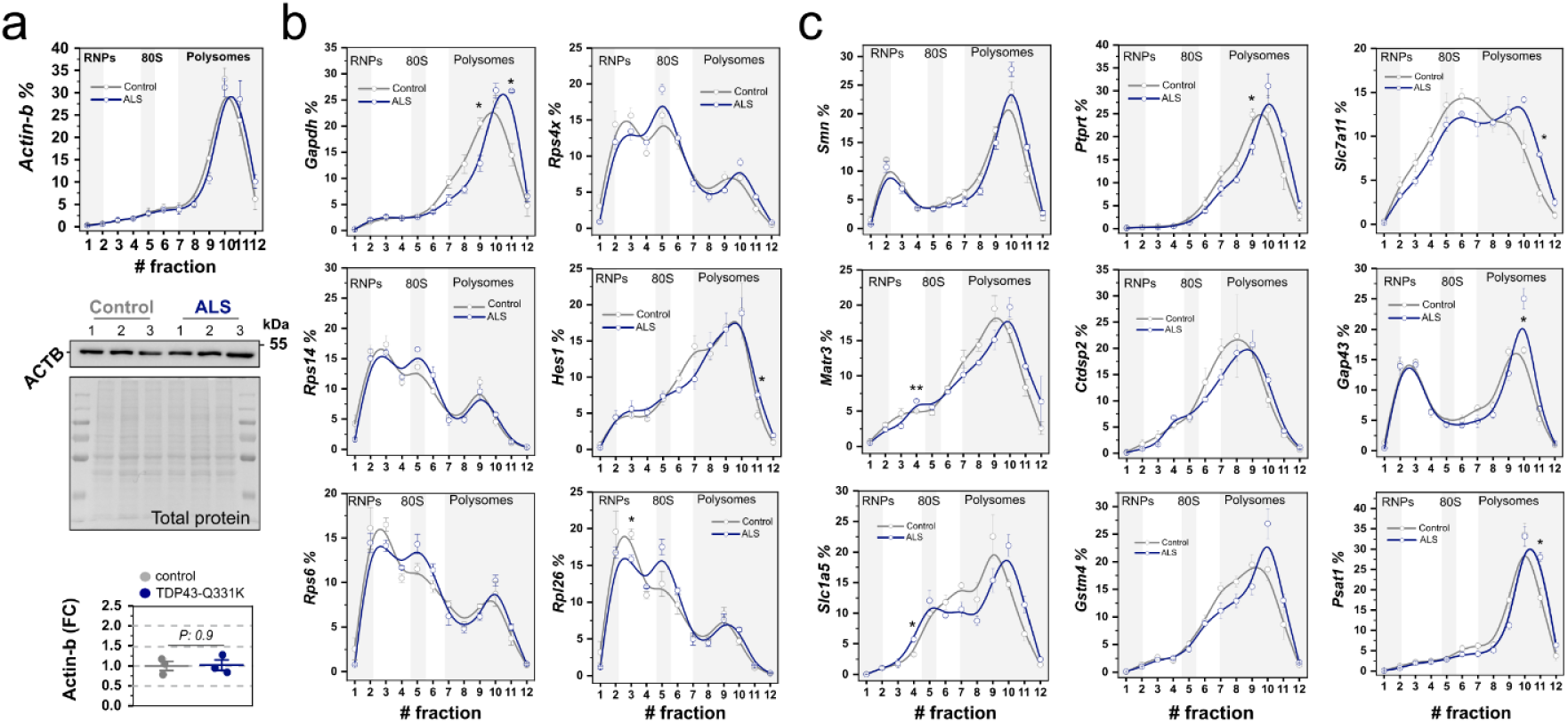
(**a**) Actin-b mRNA shows no variations at mRNA or protein levels in control and ALS spinal cords and cortices. Upper panel: relative co-sedimentation profile of Actin-b mRNA along the sucrose gradient fractions. Middle panel: protein levels of ACTB obtained using western blot analysis in n=3 biologically independent replicates. Bottom panel: relative changes of ACTB protein expression. In the upper and bottom panels, data are mean ± SEM of n=3 biologically independent replicates. Differences were assessed with two-tailed T-test. The p-value is reported in the figure. (**b-c**) Relative co-sedimentation profile of TDP-43 non-targets (b) and TDP-43 target (c) mRNAs out of the 21 transcripts selected for *in vivo* analysis. Data are mean ± SEM among n=3 biologically independent replicates. Differences were assessed with two-tailed T-test (P-value: *< 0.05, **<0.01).

## SUPPLEMENTARY Tables

**Supplementary Table 1.** List of antibodies.

| Protein | Company & code | Dilution |
| --- | --- | --- |
| ACTB | SantaCruz<br>sc69879 | WB 1:1000<br>Dot blot 1:1000 |
| ALEXA 488<br>anti-mouse | Invitrogen<br>A-11001 | IF 1: 1000 |
| ATTO 594<br>anti-mouse | Sigma<br>76085 | IF 1: 1000 |
| FUS | Novus Biologicals<br>NB 100-565 | WB: 1:7000<br>Dot blot 1:5000 |
| GAPDH | SantaCruz<br>sc-166545 | WB 1:1000<br>IF 1:200 |
| HRP<br>goat anti-mouse | Thermo Scientific<br>31430 | WB 1:1000-3000<br>Sub fract 1:1000-3000<br>Dot blot 1:1000 |
| HRP<br>goat anti-rabbit | Thermo Scientific<br>31460 | WB 1:1000-3000<br>Sub fract 1:1000-3000<br>Dot blot 1:1000 |
| HUD | SantaCruz<br>sc-2899 | WB 1:1000<br>Dot blot 1:200 |
| MATR3 | Bethyl Laboratories<br>A300-591A | WB 1:1000 |
| PABP | Abcam<br>ab21060 | WB 1:2000<br>Sub fract 1:1000<br>Dot blot 1:1000 |
| PUROMYCIN | Millipore<br>MABE343 | WB 1:5000<br>IF 1:4000 |
| RPL26 | Abcam<br>ab59567 | WB 1:1000<br>Sub fract 1:2000<br>Dot blot 1:1000 |
| RPL29 | Thermo Scientific<br>PA5-27545 | WB 1:1000 |
| RPS6 | Cell Signalling<br>2217 | WB 1:1000<br>Sub fract 1:1000<br>Dot blot 1:1000 |
| RRBP1 | Genetek<br>GT5610 | WB 1:3000<br>IF 1:125 |
| SMN | BD Transduction<br>Laboratories<br>610646 | WB 1:1000 |
| TDP43 | Proteintech<br>10782-2AP | WB 1:1000<br>Sub fract 1:1000<br>Dot blot 1:1000 |
| VIM | Genetek<br>GT812 | WB 1:3000<br>IF 1:250 |

**Supplementary Table 2.**
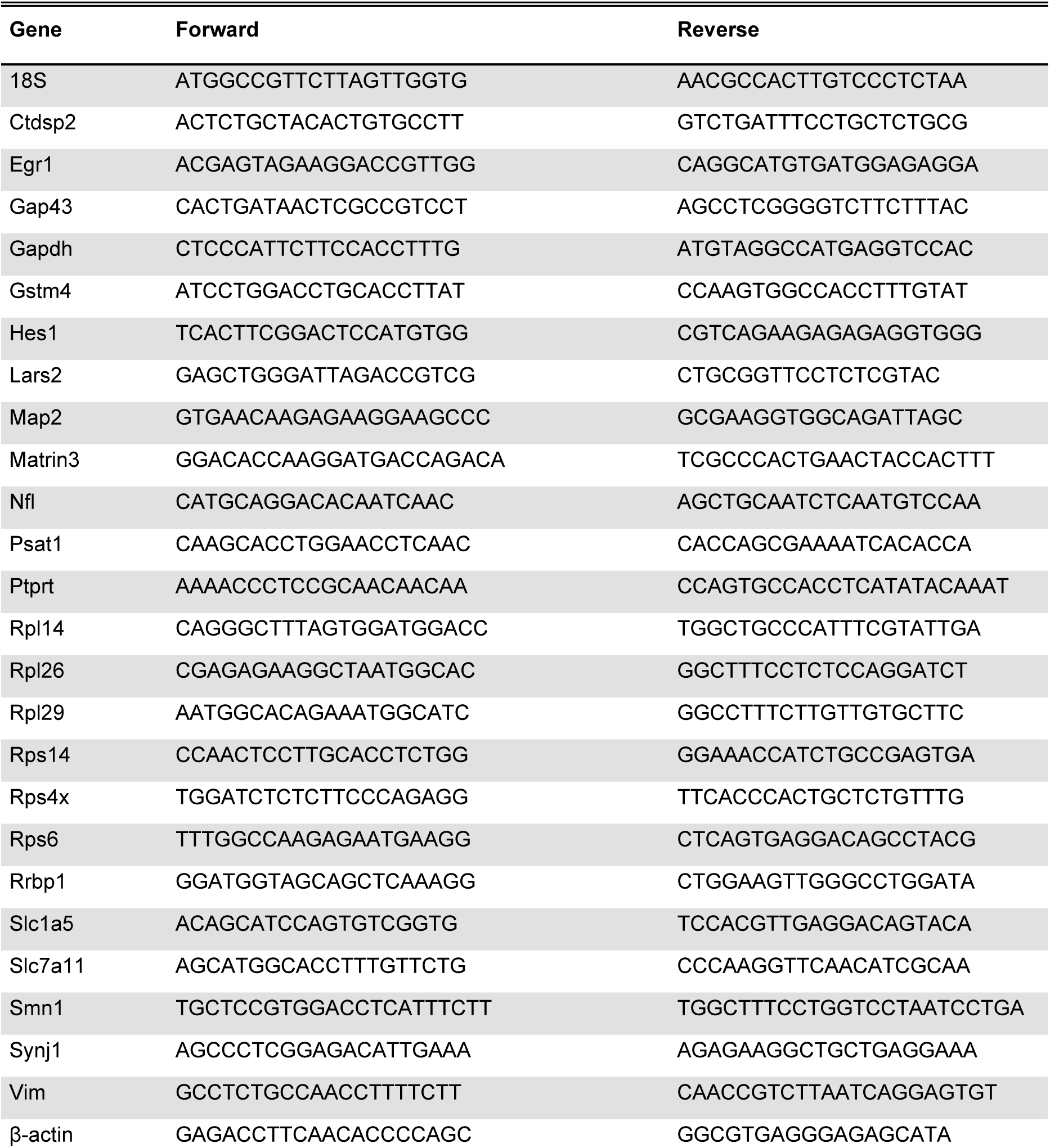
List of primers for qPCR and PCR.

## Notes

### Summary of Updates

All figures, results and discussion revised and updated

