## Supplementary File 1 for "Disruption of ribosome dynamics and mRNA homeostasis triggers a cascading imbalance in protein synthesis in models of Amyotrophic Lateral Sclerosis"

### Panel S1a

TDP-43 short exposure

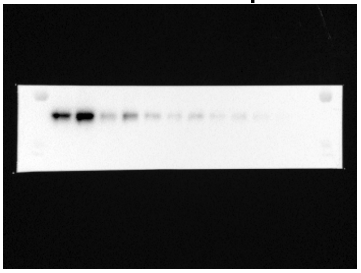

TDP-43 long exposure

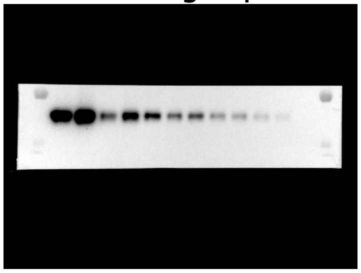

RPS6

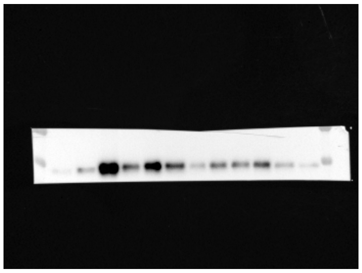

RPL29

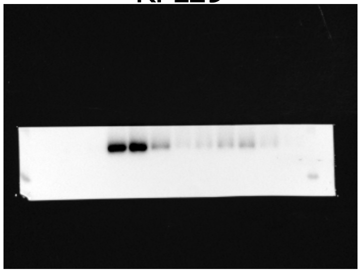

FUS short exposure

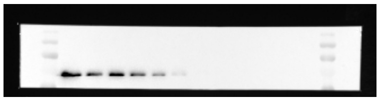

FUS long exposure

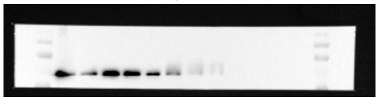

### Panel S1b

TDP-43 short exposure

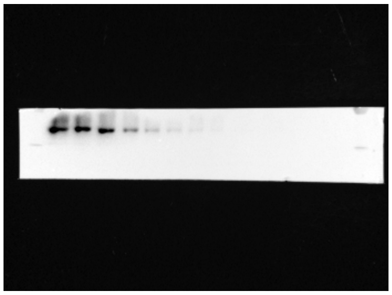

TDP-43 long exposure

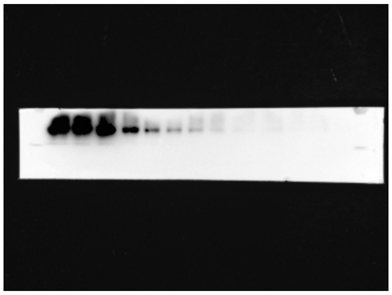

RPS6

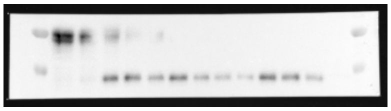

RPL26

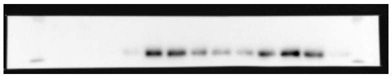

FUS short exposure

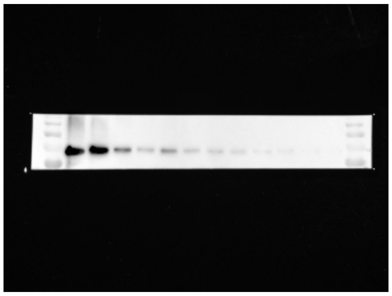

FUS long exposure

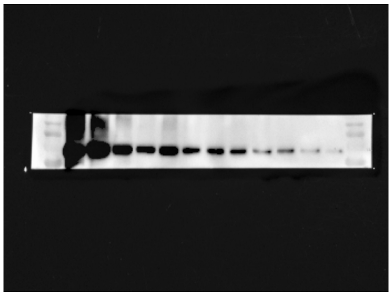

#### Panel S1c

TDP43

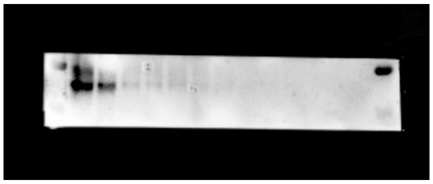

RPS6

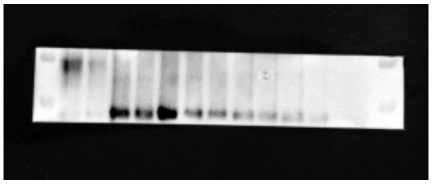

RPL26

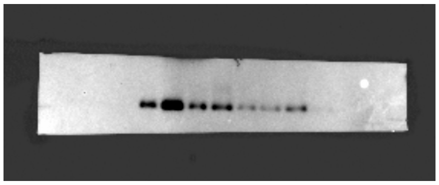

FMRP

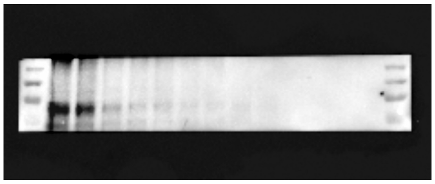

FUS

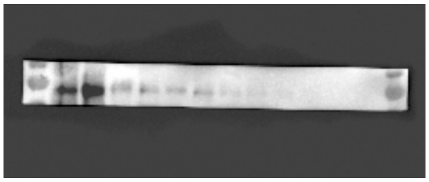

HUD

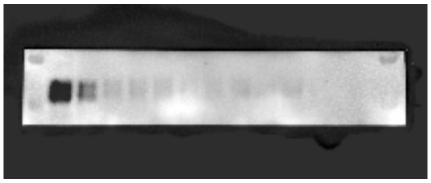

#### Panel S1d

TDP43

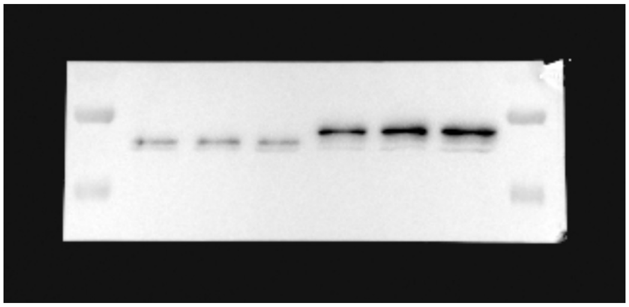

Ponceau

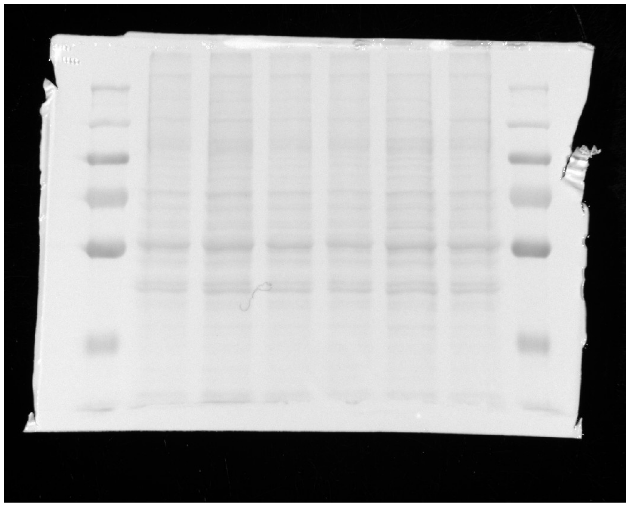

#### Panel S1e

TDP-43

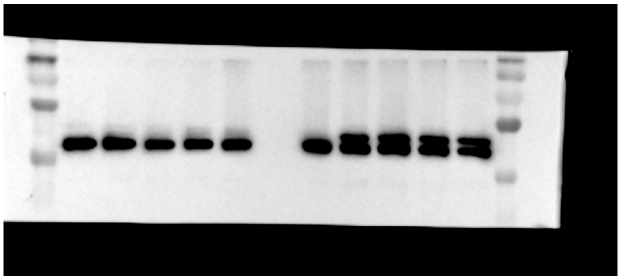

Ponceau

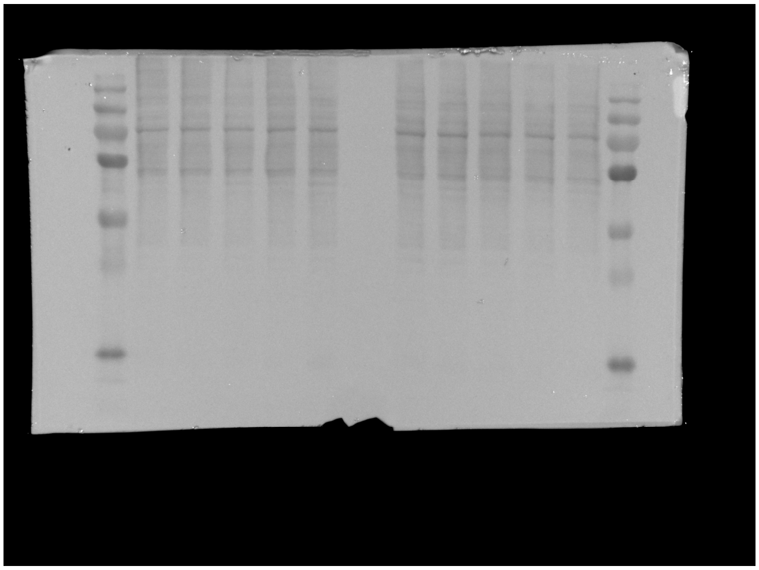
