## Supplementary figures and images for "Disruption of ribosome dynamics and mRNA homeostasis triggers a cascading imbalance in protein synthesis in models of Amyotrophic Lateral Sclerosis"

### Supplementary File 2

#### Panel 1c

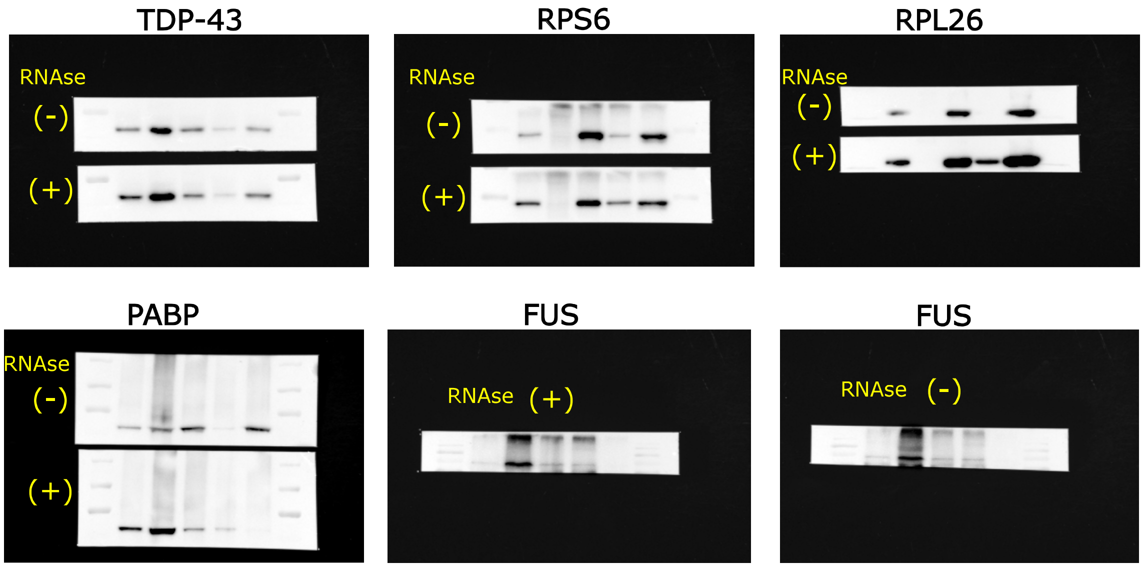


#### Panel 1e (CTRL)

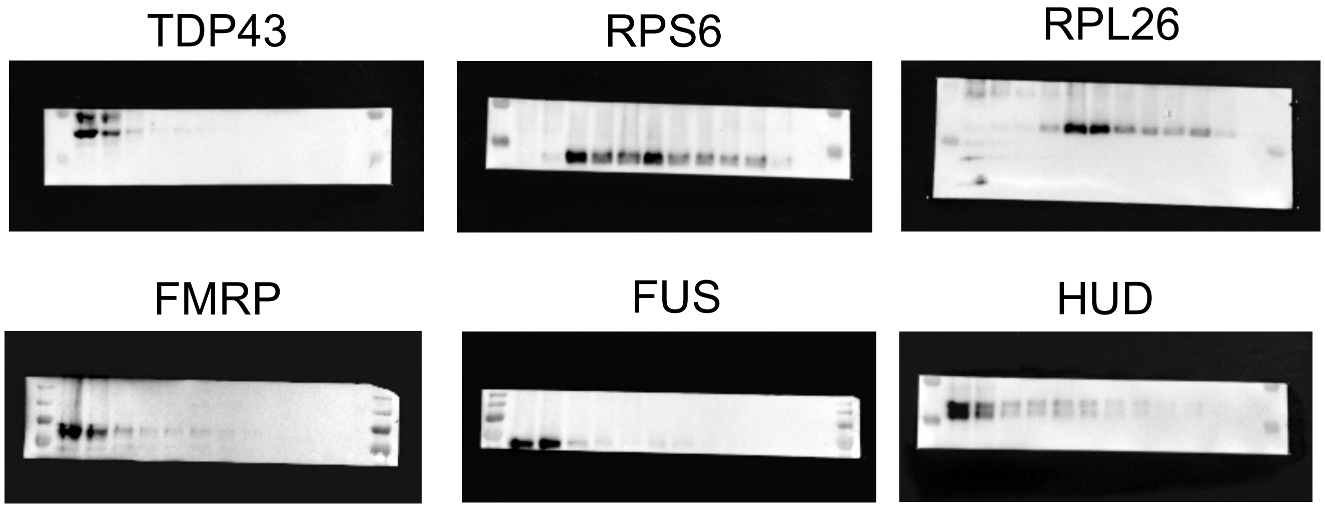


#### Panel 1e (Q331K)

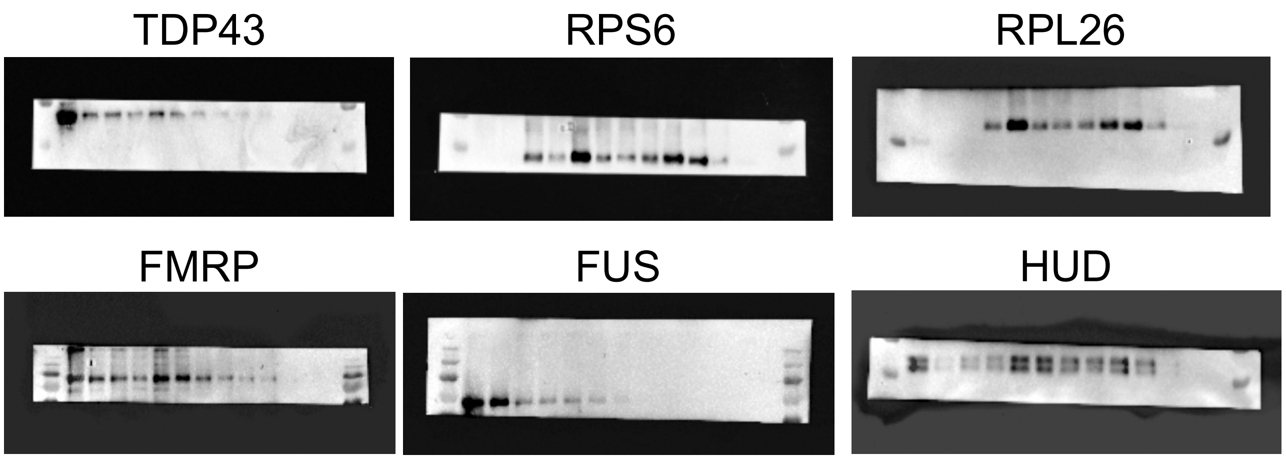

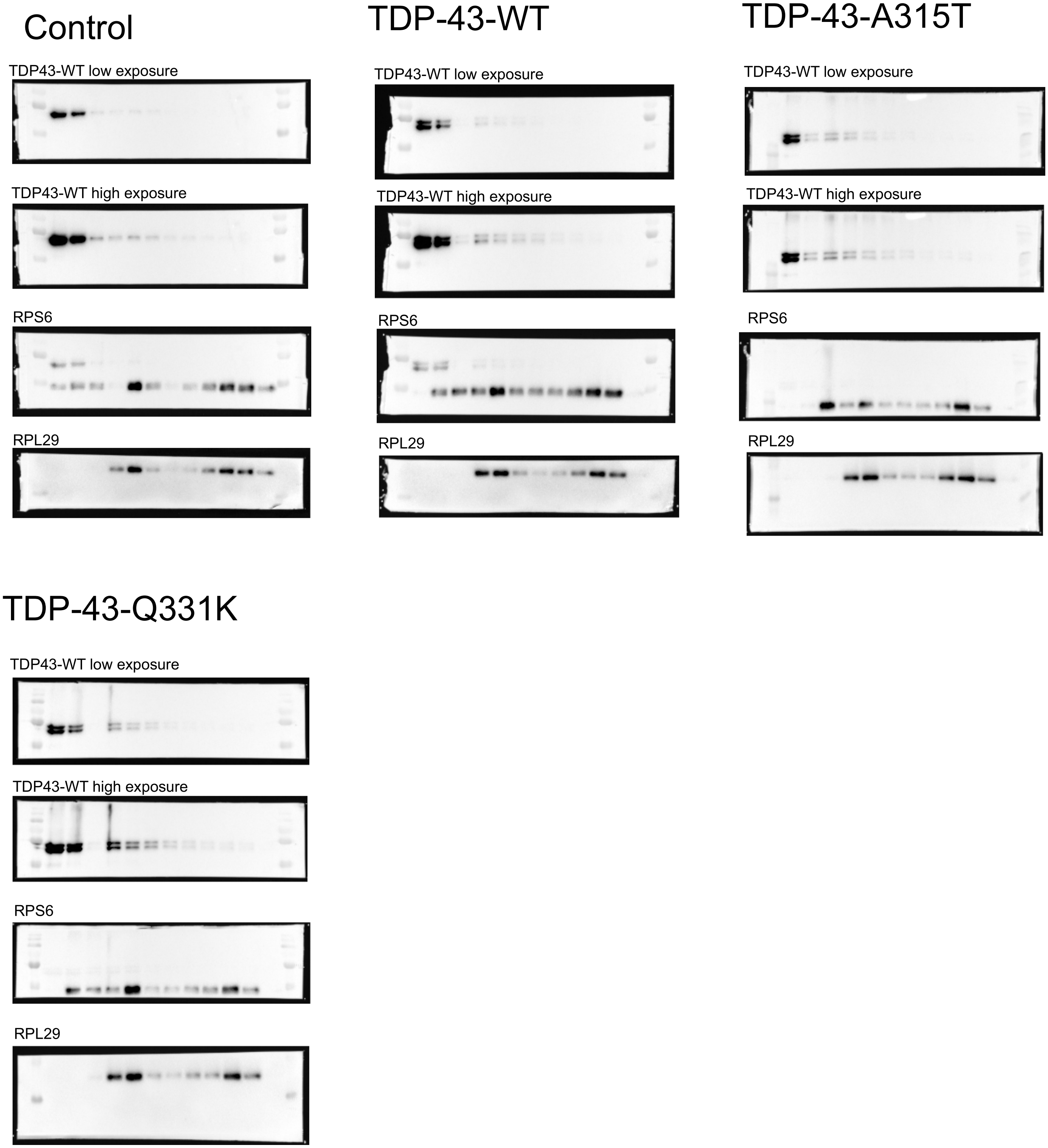

### Supplementary File 3

##### Panel S2c

NSC-34

RPS6

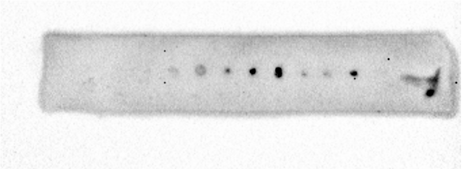

Actn b

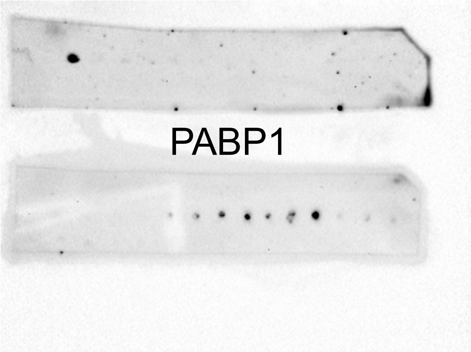

PABP1

PCNs

RPS6

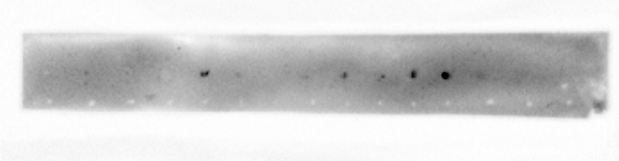

Actn b

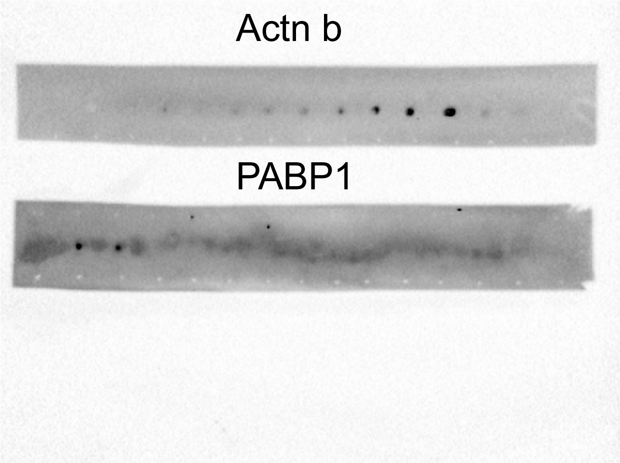

PABP1

##### Panel S2d

TDP-43

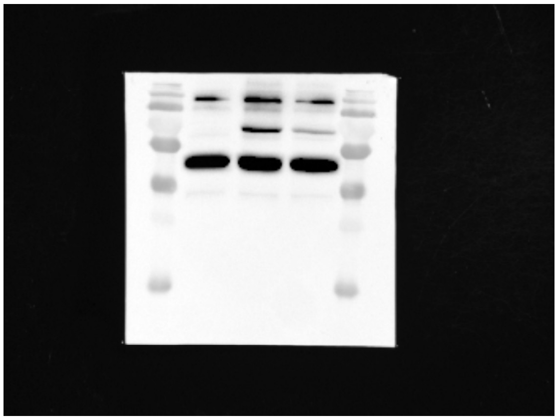

TRFP

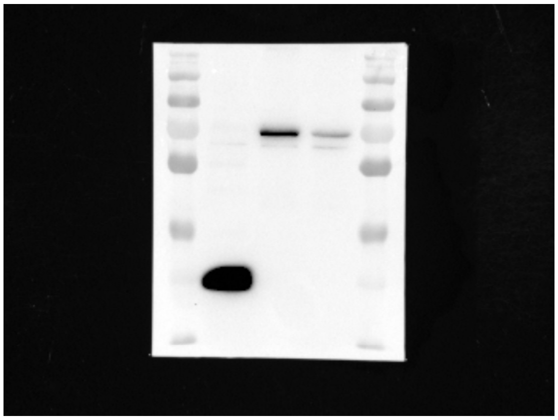

### Supplementary File 4

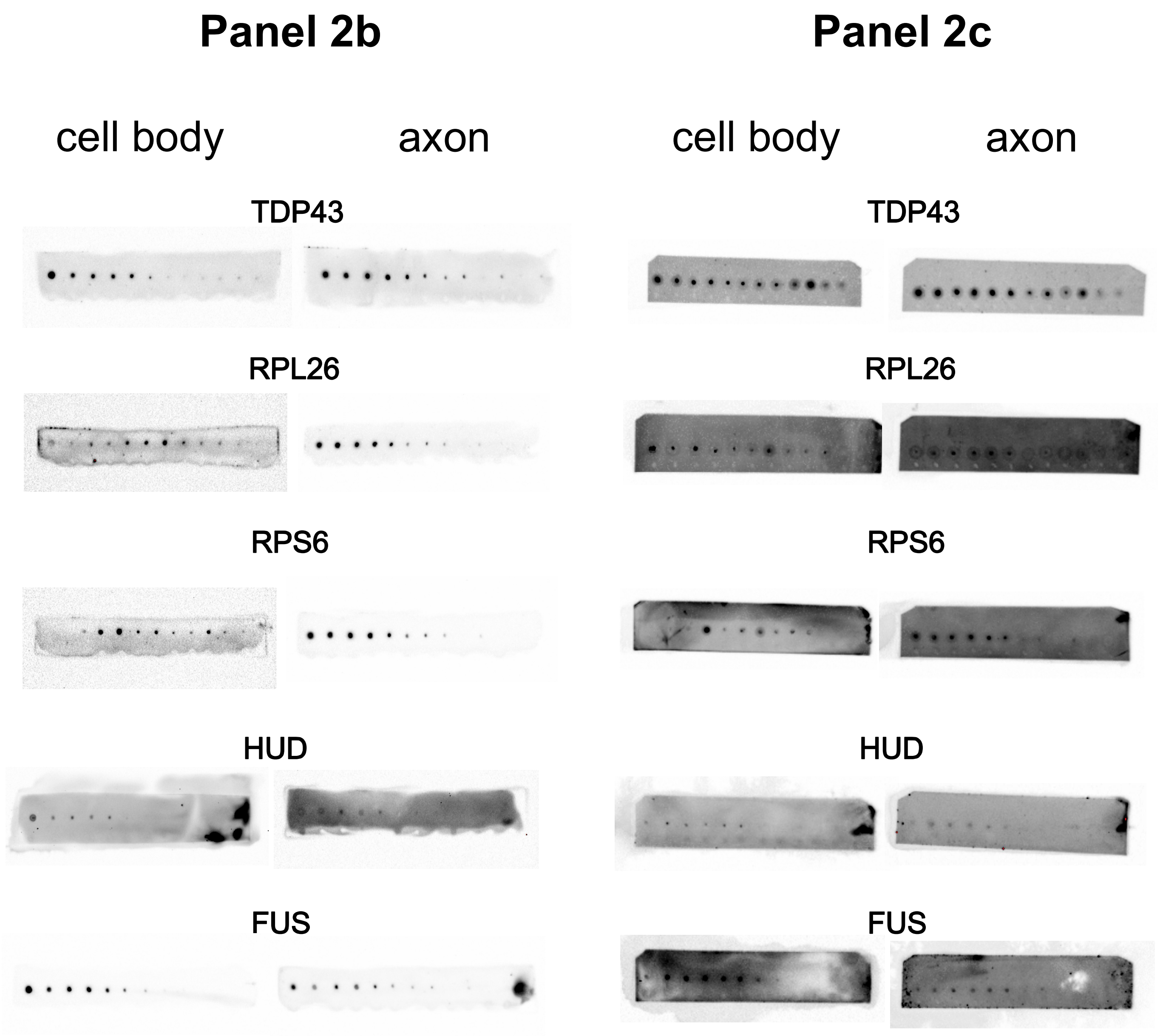

### Supplementary File 6

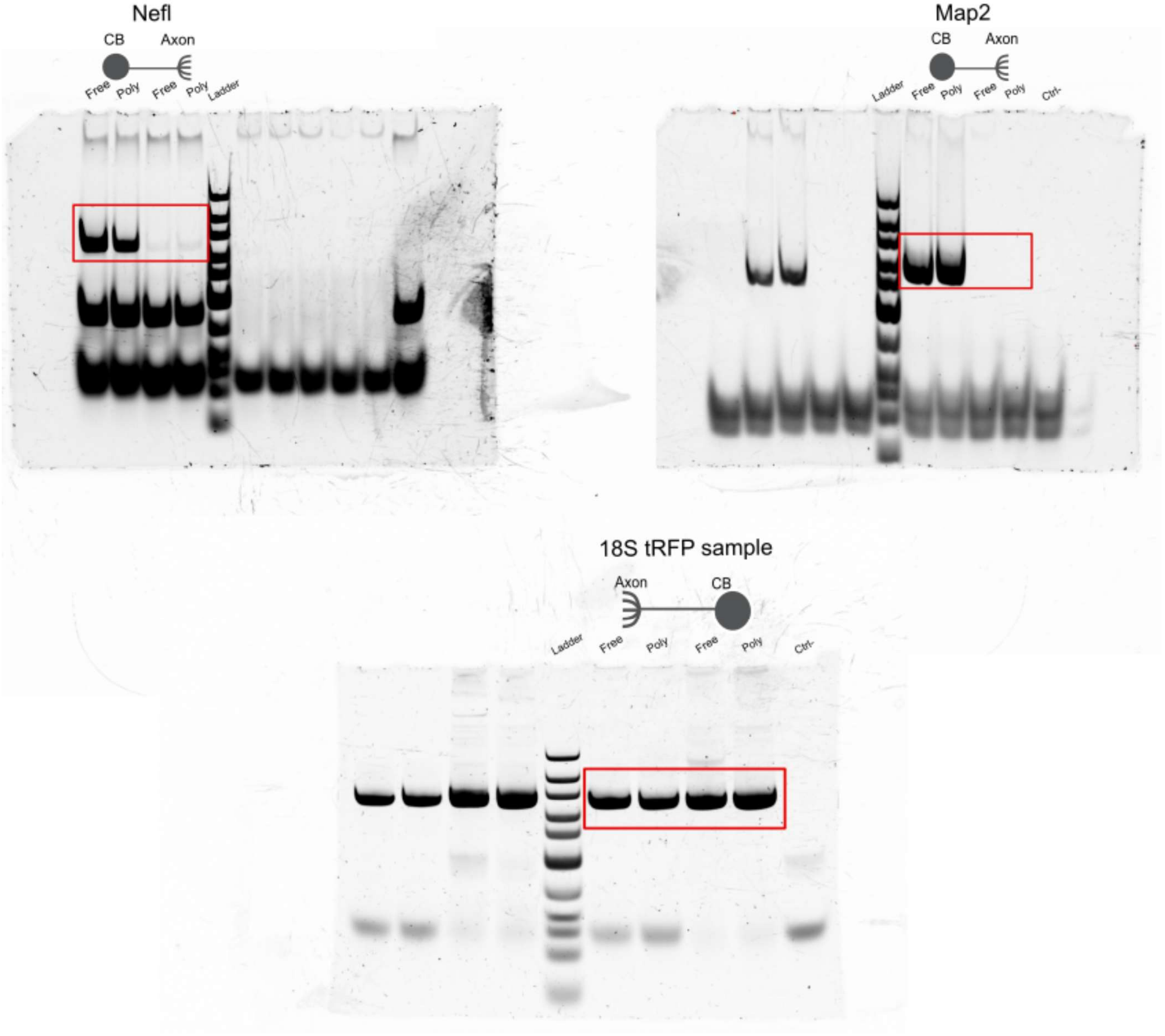

### Supplementary File 8

ACTB

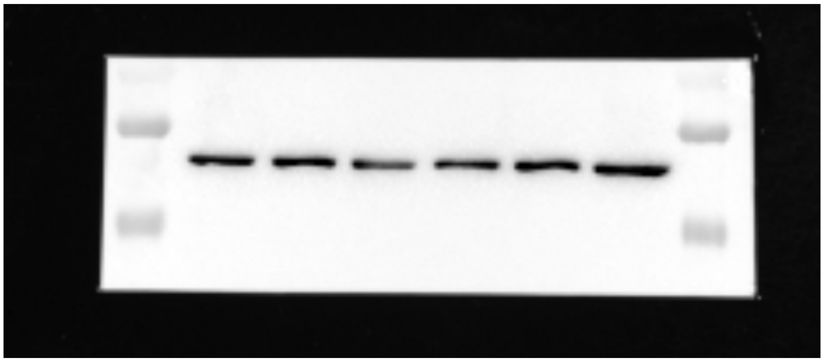

Ponceau

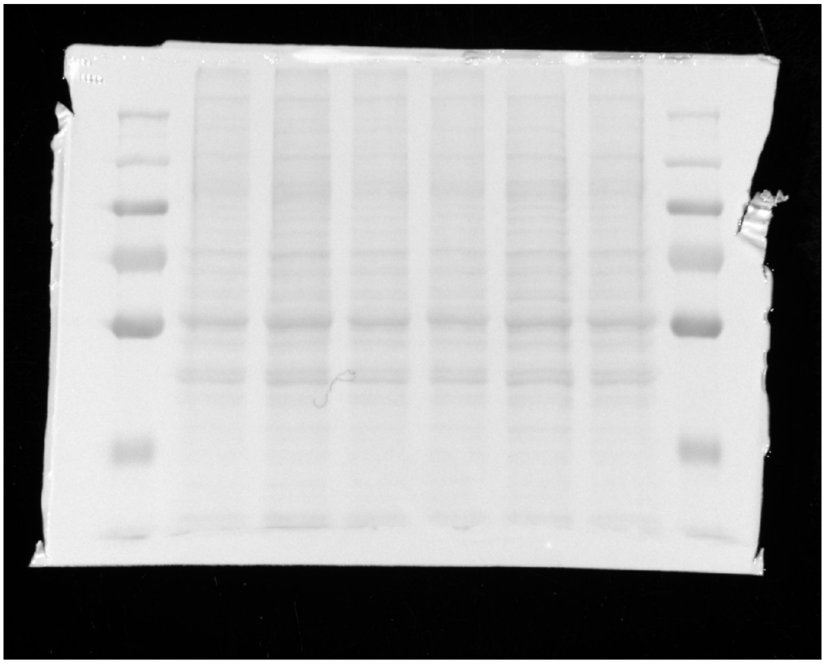
